# Identification of two novel DnaJ chaperone family proteins as modifiers of Huntingtin aggregates

**DOI:** 10.1101/2023.10.31.564873

**Authors:** Ankita Deo, Rishita Ghosh, Snehal Ahire, Amitabha Majumdar, Tania Bose

## Abstract

Huntington’s disease (HD) is a rare neurodegenerative disease. It is caused due to aggregation of Huntingtin (HTT) protein containing Q repeats more than 40. Similar protein aggregation is also a hallmark of several other neurodegenerative diseases related to loss of cognitive function. In search of modifiers of HTT aggregation, we have screened putative chaperone proteins from *Drosophila* in both fly and yeast model of HD. DnaJ chaperones were screened by evaluating HTT protein aggregation related phenotypes using growth assays for studying the growth rate of the cells, imaging studies, and gel-based approaches like semi-denaturing detergent agarose gel electrophoresis (SDD-AGE). Our screening led us to categorize several proteins as suppressors and enhancers of the HTT associated phenotypes. Out of the 40 chaperones and co-chaperones, two chaperones that came up strikingly were CG5001 and P58IPK. Protein aggregation was found to be reduced in both S2 cells and *Drosophila* transgenic lines with HTT103Q in presence of these class of chaperones. As these DnaJ chaperones have protein sequence similarity across species, these might be used as possible tools to combat the effects of neurodegenerative diseases, as evidenced specifically in Huntington’s disease and Amyotrophic Lateral Sclerosis (ALS).

## Introduction

Neurodegenerative diseases are a class of neurological disorders characterized by gradual dysfunctions associated with loss of neurons. Protein deposition in the human brain is a phenomenon that is fundamental in most neurodegenerative disorders [1]. A slow continuous loss of neural cells causes neurodegenerative diseases leading to nervous system dysfunction [2]. These diseases have diverse pathophysiologies ranging from cognitive impairment to difficulties in performing day-to-day tasks.

Our study focuses on protein aggregations in Huntington’s disease supported by a few observations regarding toxicity and its recovery in the background of Amyotrophic Lateral Sclerosis (ALS).

ALS involves a continuous degradation of motor neurons. Thus, the motor neurons become susceptible to oxidative stress along with stress granule formation and mitochondrial damage [3]. TDP-43 (Tar DNA binding protein 43) [4] and FUS (Fused in Sarcoma) [5–7] are two proteins that link ALS to oxidative stress and stress granule formation.

Huntington’s disease is a neurodegenerative condition resulting from an expansion of CAG repeats on the gene coding for Huntingtin protein on chromosome 4p16.3 [8]. While the striking feature of the disease is the involuntary movements of the muscles also known as ‘Chorea’, the other symptoms include psychological abnormalities and neuronal loss in the brain [9]. Expansion of CAG repeats results in addition of Glutamine (Q) stretches in the exon 1 of the Huntingtin gene and when it exceeds more than 40, it increases the occurrence of the disease [10]. Huntingtin has many repeated caspase cleaving sites. In Huntington’s disease, cleavage of mutant HTT releases N-terminal fragment with increased toxicity and aggregation because of the presence of the expanded poly-Q tract. This leads to the onset of neurodegeneration in Huntington’s disease [11]. The Expanded poly Q stretches cause the HTT protein to misfold and form aggregates causing the disease. One way to alleviate the phenotypes associated with HD is to decrease the aggregate load.

It has been observed that in yeasts, the aggregates in Huntington are correlated to the length of the poly Q repeats, suggesting that aggregation and pathogenicity are dependent on the increase of the poly Q repeat and could be regulated by the expression of chaperone proteins [12].

Chaperone proteins play a crucial role in helping disassemble protein aggregates or target proteins for degradation. Thus, they act as a backbone of protein quality control checks. Simultaneously, these cellular chaperones regulate the folding and maturation of newly synthesized as well as partially folded proteins within the cell. They also help resolve the aggregated misfolded proteins [13].

Heat shock proteins (Hsps) are specialized chaperones with the ability to control proteostatic stress [14]. Hsps can be characterized based on their molecular weight, structure, and functions. Some examples are Hsp10, Hsp40, Hsp60, Hsp70, Hsp90, and Hsp110 [15]. Small molecular weight Hsps can be classified based on the pattern of their expression, which may be either, ubiquitous or tissue-specific [16].

Here, we have generated a library of putative Hsp40 and related chaperones from *Drosophila* for expression in yeast and used this library to initially screen a HD model in yeast to look for modifiers of Huntington disease related toxicity. Although, there have been reports of using the DnaJ library in yeasts, to study the Huntington aggregates, we undertook this study by using an extensive overexpression screen to complement the previous studies in a quest to exploit the disaggregase properties of additional DnaJ domain proteins. We further tested these genes in the S2 cell line and fly hemocytes to check if the phenotypic traits match those effects found in yeasts.

Work done on Hsp40 chaperones in yeast Huntington models report overexpression of the yeast Hsp40 chaperone, like sis1, decreases polyglutamine aggregate size and toxicity [17]. Overexpression of Ydj1, another yeast Hsp40 chaperone was found to modulate the physical properties of the Huntington diseased (HD) exon 1 aggregates by suppressing the formation of SDS insoluble HD aggregates [18]. The chaperones thus help in protein quality control [19] and help yeast combat stress induced by heat shock or other parameters [20].

Hsp40s have a J domain, making them members of the DnaJ domain chaperone family. J domain of DnaJ chaperones is a 70 amino acid long region conserved across prokaryotic and eukaryotic chaperones [19] J protein activates the Hsp70 ATPase in the chaperone cycle [21]. It transfers the substrate protein to the substrate binding domain of Hsp70. This leads to the folding of the substrate protein. Then J protein dissociates from Hsp70 and the substrate protein is released. J proteins have three structural classes across species with a compact helical J-domain present in all 3 of these classes [19]. The first class is the Type I-J proteins DnaJAs. They are distinctive because of cysteine-rich zinc finger domain. There is a glycine/phenylalanine-rich region that links the J-domain to the cysteine-rich zinc finger domain [22,23]. The second class is the Type II-J proteins DnaJBs. They do not contain the cysteine-rich region but they have a J-domain connected by glycine/phenylalanine-rich region to the C-terminal domain. This class is especially known to aid in protein disassembly [24].

The third class is Type III-J proteins DnaJCs which have only the J domain stretch. They are very varied in carrying out cellular functions that differ from typical Hsp70 co-chaperone functions [25,26].

DnaJ proteins have multiple protein domains including ubiquitin interacting motifs or clathrin binding domains which assist in protein misfoldings in cells in ER stress mediated diseases [27]. They also contain mitochondrial leader sequences or ER signal peptides targeting them to specific cell organelles that help them combat the misfolding stress due to neurodegenerative diseases [27].

Another function of DnaJ class chaperones is to prevent protein aggregation and solubilize the aggregates when combined with Hsp70 and nucleotide exchange factors [27]. A classic interplay of Hsp40 and Hsp70 plays a crucial role in protein quality control in neurodegenerative diseases. Hsp40 chaperones interact with Hsp70 protein family and function as cofactors for different substrates. In some cases, Hsp70 interacts with Hsp40 to act as co-chaperones for translating ribosomes [28,29]. *Drosophila* has both the heat shock inducible Hsp70s and the constitutively expressing Hsc70s [29,30]. Taking cues from here, we set about to do a screen to look for enhancers and suppressors initially in a yeast model using the 35 Hsp40 proteins, 3 Hsc70 (Hsc70-1, Hsc70-4, and Hsc70cB) and 2 Hsp70 (Hsp70Aa, Hsp70Bb) due to redundancy in their sequence with others.

According to the prediction models, we have chosen the chaperones with a role in protein folding activity. The functional activity of their human homologs suggests the same [30,31].

We screened 35 DnaJ domain chaperones from *Drosophila* which are: Mrj, CG5001, DnaJ60, CG6693, CG2911, JdpC, CG4164, CG9828, CG2887, CG30156, CG12020, CG10565, Wus, Tpr2, Csp, CG7394, CG8476, CG7130, CG7872, Sec63, Hsc20, P58IPK, Droj2, CG43322, CG7133, CG14650, CG7556, CG32641, CG10375, CG11035, CG3061, CG17187, DnaJ1, CG7387 and CG8531 in the Huntington model of yeast. These chaperones mostly function in protein unfolding activities [30–32].

For example, CG5001 has a predicted chaperone binding activity and unfolded protein binding activity [30]. It is orthologous to several human genes including DnaJB5 (DnaJ heat shock protein family (Hsp40) member B5) and is located in the cytosol [33,34]. DnaJB5 acts as a novel biomarker and a possible target for cervical cancer treatment. Knock down of DnaJB5 decreases proliferation and increases apoptosis of SiHa cells, which are cells of uterine tissue with squamous cell carcinoma [34,35].

P58IPK is predicted to enable chaperone binding activity and misfolded protein binding activity. It is involved in protein folding in the endoplasmic reticulum and is orthologous to the human DnaJC3 (DnaJ heat shock protein family (Hsp40) member C3). DnaJC3 deficiency in mice causes pancreatic β cell loss and diabetes. Loss-of-function mutations in DnaJC3 result in multisystemic neurodegeneration and early-onset diabetes [35,36]. It is also known for negative regulation of ER stress induced eIF2α phosphorylation [36,37].

CG7556 is expressed in adult fly head, embryonic/larval salivary gland, and testis. It is orthologous to human DnaJC1 (DnaJ heat shock protein family (Hsp40) member C1). DnaJC1 regulates protein secretion and is located in the endomembrane system [37,38] and endoplasmic reticulum.

CG7133 is a protein coding gene from *Drosophila*. It has one annotated transcript and one polypeptide.

CG11035 is predicted to be involved in brain development and regulation of mitochondrial ATP synthesis coupled proton transport. It is orthologous to human DnaJC30 (DnaJ heat shock protein family (Hsp40) member C30). DnaJC30 is involved in Williams-Beuren syndrome and nuclear type mitochondrial complex I deficiency. It is located in the mitochondrial inner membrane [38,39].

CG8531 is predicted to enable ATPase activator activity. It is orthologous to human DnaJC11 (DnaJ heat shock protein family (Hsp40) member C11). DnaJC11 is involved in cristae formation in the mitochondrion [39–41]. CG43322 is predicted to be involved in the Golgi organization, it is orthologous to human DnaJC28 (DnaJ heat shock protein family (Hsp40) member C28) and is present in the Golgi transport complex.

CG12020 is predicted to enable chaperone binding activity and unfolded protein binding activity. It is involved in chaperone cofactor-dependent protein refolding. It is orthologous to human DnaJB13 (DnaJ heat shock protein family (Hsp40) member B13). It is present in the axoneme, cytosol, and sperm flagellum.

Apart from these 8 chaperones, others like Mrj and Droj2 have also been studied. Mrj has chaperone binding activity, unfolded protein binding activity, and chaperone cofactor-dependent protein refolding. It is orthologous to several human genes including DnaJB6 (DnaJ heat shock protein family (Hsp40) member B6). DnaJB6 is a general conditioner of toxicity and solubility of multiple ALS/FTD (amyotrophic lateral sclerosis and frontotemporal dementia) linked RNA binding proteins. It co-phase separates with and alters the behavior of FUS (fused in sarcoma) condensates by preventing their fibrillization [41,42].

Droj2 enables chaperone binding activity. It is involved in protein refolding. It is orthologous to the human DnaJA1 (DnaJ heat shock protein family (Hsp40) member A1) and DnaJA4 (DnaJ heat shock protein family (Hsp40) member A4). Knockout of DnaJB6 in Huntington’s model of HEK293 cells results in an increase in 74Q HTT aggregation whereas knockout of DnaJA1 results in a decrease in aggregation. DnaJB6 and DnaJA1 modulate poly Q toxicity in opposite manners [42,43]. Here, Hsp40, and related chaperones from *Drosophila* are screened for their expression in HD model of yeast.

Huntington’s disease model is developed in *Saccharomyces cerevisiae* by expressing 25Q and 103Q constructs with 25 and 103 CAG repeats respectively [43]. These constructs are under the influence of galactose inducible promoters. A simple 2% galactose induction allows the protein with the Q repeats to be induced in the cell. Similarly, mutants of ALS were made by introducing galactose inducible plasmids pRS426 Gal-TDP-43-GFP [44] and pAG416 Gal-FUS-YFP [45] which are TDP43 and FUS plasmids in host *S. cerevisiae*.

25Q serves as control whereas the mutant 103Q expresses the Huntington phenotype [43]. 25Q grows normally and the protein is diffused in the cell cytoplasm. We have chosen the HTT mutant with 103 CAG repeats such that it shows growth defects at permissive temperature associated with protein aggregation, but is not lethal. These features have been characterized via growth assays, imaging, and electrophoretic techniques. Protein aggregation characteristic of the disease intrigued us to check if the chaperones would play a certain role in the disease management.

Addressing the pathogenic conditions of Huntington disease prompted us to look for potential Hsp40 DnaJ chaperones, which would either aggravate or ameliorate the slow growth and protein aggregation phenotype. This led us to perform an overexpression screen of DnaJ domain proteins of flies and expressing it in the yeast Huntington 103Q strain. S2 is the most widely used *Drosophila* cell line with mesodermal traits and are derived from hemocytes [46]. S2 cell line was used to study the functional properties of the modifier chaperones after extensive screening in the yeast system. Further, the crosses set up using transgenic flies show clearly the decrease in pathogenic aggregates of HTT103Q in hemocytes co-expressing CG5001 and P58IPK respectively.

Since these chaperones are derived from flies, we decided to look at the functional effect of these in S2 fly cell line and HD models of *Drosophila*. The fruit fly *Drosophila melanogaster* is used extensively as an animal model in biology. A lot of cellular and molecular tools have also been developed or adapted to work with *Drosophila* in the past two decades including the complete genome sequencing. The *Drosophila* system has an advantage of combination of classical genetics and molecular and cellular techniques. This has encouraged researchers to study the pathophysiology of neurodegenerative diseases in terms of gene expression, behavior, development, and aging [46,47]. Expression of poly Q tract in *Drosophila* leads to retinal degeneration. Poly Q diseases include Spinocerebellar ataxia type 1, 2, 3, 6, 7, 17, Spinobulbar muscular atrophy, Dentatorubralpallidoluysian atrophy, and Huntington’s disease [47,48]. Small changes in the gene function are intensified in the fly’s eyes. Since eyes are non-essential for the survival of flies this can be exploited to carry out genetic and chemical screenings of poly Q modifiers on a large scale. Proteinaceous neuronal inclusions accumulate in fly brains as a result of poly Q diseases [49,50]. *Gal4-UAS* binary system can be used to manage time specific expression of target genes in neurons [51,52]. *Drosophila* model of neurodegenerative diseases have been constructed using transgene techniques. Molecular chaperones have been reported to suppress neurodegeneration in *Drosophila* [53,54].

Phenocopying the candidate genes found in yeast and expressing them in *Drosophila* S2 cell lines as well as in transgenic flies helped us further understand the pathophysiology of the diseased condition.

We have used chaperones and co-chaperones in yeasts and fly models to show that the pathogenicity in neurodegenerative diseases can be altered by selective DnaJ modifier proteins.

## Results and Discussion

### Expression of Hsp40 and Hsp70 proteins of *Drosophila* in yeast

Yeast cells are widely used to study various neurodegenerative diseases. Huntington’s disease in particular has been studied in yeasts in the quest of exploiting the translational potential of the system [55,56]. To check for toxicity in Huntington’s disease, we used the 25Q repeats as control and 103Q repeats as Huntington mutants and transformed them in yeasts with the appropriate plasmids following already published work [54,57,58]. The poly Q repeats are under the influence of a galactose inducible promoter. 25Q repeats did not show any toxic phenotypes of the Huntington’s mutation [48]. In the 103Q mutant, the growth is slowed down as evidenced from the galactose plates in serial dilution assay after 48hrs. However, growth on the dextrose plate for both 25Q and 103Q remains the same (Supplemental Fig.1A). Simultaneously, we wanted to check if the Huntington mutant shows any kind of aggregation due to the repeat units. The 25Q strain shows a diffused cytoplasmic GFP expression, but the 103Q mutant shows distinct cytoplasmic GFP punctae under confocal microscope 24 hrs post induction of the strain with galactose (Supplemental Fig.1D).

As Huntington’s disease mutant displayed slow growth and protein aggregation, we screened DnaJ domain chaperones for their effects on these phenotypes of the mutant. DnaJ proteins belong to the Hsp40 class of protein quality control chaperones. This genetically and functionally diverse class of chaperones comprises of 22 proteins expressed in yeast *Saccharomyces cerevisiae,* 49 in humans, and 43 in fly *Drosophila melanogaster*.

Members of the Hsp40 protein family play a role in numerous processes involving protein folding and refolding. Liberek et al, first reported that function of DnaJ proteins is for assisting Dnak (bacterial Hsp70) in increasing its ATPase activity. This was deciphered *in vitro* by purifying DnaJ from *E. coli* [56,59]. It is known previously that DnaJ helps Hsp70 and its DnaK in disaggregation of proteins *in vitro* [57,58,60,61]. A study by Acebron et al. [59,62] also reports DnaJ working with DnaK and nucleotide exchange factor GrpE to prevent protein aggregation. DnaJ also helps to solubilize the aggregates alone or with help of *E. coli* Hsp100 representative ClpB [59,62]. DnaJ has an essential role in almost all Hsp70 chaperone activities. DnaJ chaperones act as the first blockade against protein aggregation as Hsp40 recruits Hsp70 in stressed conditions. These studies were followed up by researchers trying to manage the toxicity caused due to protein aggregation diseases by using DnaJ chaperones. Cytosolic members of DnaJA and DnaJB subfamilies have been reported to suppress Parkin C289G aggregation in Hsp70 dependent manner [60,63]. Amyotrophic Lateral Sclerosis is another neurodegenerative and protein misfolding disease. TDP-43, FUS, and SOD1 are some of the proteins that mislocalize in neurons of ALS patients. They are also found to partition in cellular inclusions in NSC-34 motor neuron-like cells [61,64]. Overexpression of DnaJB1 and DnaJB2 have shown protective effects in ALS models as well. Overexpression of DnaJB1 and its yeast homolog Sis1 decreases TDP-43 mediated toxicity by affecting cell growth, cell shape, and ubiquitin-proteasome system inhibition [62,65]. On the other hand, overexpression of DnaJB2 improves muscle performance and motor neuron survival in mutant SOD1, carried out in *in vitro* models of ALS [63,66].

We wanted to find out if the diseased phenotype shown by the Huntington’s disease mutant would be rescued or aggravated in presence of overexpression of DnaJ domain chaperone proteins. There was a high degree of sequence similarity in the yeast Hsp40, Hsp70, and Hsp104 proteins with *Drosophila* and mammalian homologs. *Drosophila* genome has members of Hsp40 and Hsp70 families but lacks Hsp104 homologs [64,67]. *Drosophila* has 39 Hsp40/DnaJ domain protein genes [30], 7 Hsp70, and constitutively expressing six of the Hsc70 chaperones [28,65]. We chose 35 *Drosophila* Hsp40 proteins, 2 Hsp70 proteins and 3 Hsc70 proteins for screening based on their predicted and *in situ* functions [30,31]. Huntington’s disease is a protein misfolding and aggregation disorder. DnaJ co-chaperones transfer the misfolded proteins to Hsp70 which plays an important role in the management of these protein aggregates [42,43]. In this study, Hsp40/DnaJ proteins were selected based on their ability to bind unfolded proteins, as reported for their human homolog counterpart or from predicted modeling studies in *Drosophila*. Each of the Hsp70 proteins were selected based on their sequence variability from each other.

For the expression of Hsp40, Hsp70, and Hsc70 in yeast cells, Topo-D-Entr vectors were cloned in destination vector pAG424-GAL-ccdB-HA with 3X HA tag at the C terminal [33]. Cloning of these chaperones was done with the help of destination vectors pUASg-HA for Hsp40 and pUASt-ccdB-FLAG (pTWF) for Hsp70 and Hsc70 in S2 cells.

### Enhancers and Suppressors classification following growth assays in Huntington’s mutant in yeast

DnaJ/Hsp40 class actively works against disease toxicity in models of *E. coli, Drosophila melanogaster*, *Saccharomyces cerevisiae, Mus musculus*, human cell line HEK293 [66–69]. A study from *Drosophila* sheds light on Hsp40 and its co-chaperone Hsp70 *in vitro* suppressing the formation of large detergent insoluble poly Q HTT aggregates. The activity of Hsp40 and Hsp70 results in accumulation of detergent soluble inclusions. Thus, these chaperones have an ability to shield toxic forms of poly Q proteins and direct them into non-toxic aggregates [18] DnaJ1/HDJ1 which is *Drosophila* ortholog of human HDJ1 suppresses neurotoxicity in fly model of Huntington’s disease [68,70]. Full length DnaJB14 and DnaJB12 both have protective properties against mutant FUS aggregation in an Hsp70 dependent manner. Overexpression of *Drosophila* DnaJ1 which encodes a protein homologous to human chaperone HDJ1 suppresses poly Q toxicity in fly model of spinocerebellar ataxia type-1[69,71]. Yeast strains exhibiting prion domains are also used to study effects of chaperones on protein aggregations. Ssb1, Ssa1, and Ydj1 are three yeast chaperones that cure self-propagating protein determinants on overexpression in Sup35C prion strain [70,72]. *Drosophila* Hsp40 chaperone Mrj and its mammalian homolog interact with Hsp70 proteins and prevents aggregation of pathogenic Huntingtin [28,65]. Poly Q toxicity in *Saccharomyces cerevisiae* is mediated by physical crosstalk within a network of cellular Poly Q and prion like proteins and might be caused by favorable sequestration of crucial nucleolar and mitochondrial proteins [71,73]. Zuotin, an yeast DnaJ chaperone binds RNA and associates with the ribosome, and functions with another molecular chaperone SSB [72,74]. Zuotin is homologous to human chaperone DnaJC2 and *Drosophila* chaperone CG10565. Since Huntington’s disease is also translationally controlled [54,73], we hypothesize that these chaperones may possibly play a role in reducing the translational [57] and nucleolar defects [74–76] seen in HD.

Taking cues from these findings, we screened the 40 DnaJ domain chaperones and co-chaperones from *Drosophila* for their effects on mutation of Huntington’s disease. Huntington’s mutation results in slow growth and protein aggregation at a cellular level in yeast. The DnaJ domain chaperones and co-chaperones from *Drosophila* were screened in the yeast background.

In order to understand the effect of chaperones on growth of yeast in the Huntington’s background, *Saccharomyces cerevisiae* haploid yeast was first transformed with Huntington mutant 103Q construct [58,76]. This construct had a Ura marker and a GFP tag at the C terminal. Chaperones when transformed with mutant HTT can either enhance or suppress the growth and protein aggregation effects of the mutation. This was determined by 3 independent experiments. Initially, we checked by a serial dilution assay where mutant 103Q was transformed individually with these 40 chaperones. The mutant strain was transformed with DnaJ domain chaperones and co-chaperones with an HA epitope tag at the C terminal. The transformed colonies were selected on media lacking Uracil and Tryptophan (SD Trp^-^ Ura^-^). These colonies were then serially diluted and spotted onto SD Trp^-^ Ura^-^ and Gal Trp^-^ Ura^-^ plates. Plates with Dextrose (SD) served as control, whereas galactose plates were observed for growth, as the HTT plasmids were under the effect of a galactose inducible promoter. Those chaperones that made the Huntington mutant grow slower in comparison to the Huntington mutant with empty vector Luciferase in Gal Trp^-^ Ura^-^ media enhanced the effect of the mutation, hence these were collectively referred to as enhancers. The chaperones that made the HTT mutant grow better than empty vector Luciferase on the Gal Trp^-^ Ura^-^ plate were termed as suppressors of the mutation, since, they were suppressing the slow growth phenotype of the mutant. Out of the 35 DnaJ domain chaperones P58IPK, CG5001, CG7556, Sec63, CG 2911, CG7133 suppress the mutation and others like CG12020, CG8531, CG9828, CG43322, CG11035, CG7387 enhance the mutation. All these chaperones had a role in protein folding activity as suggested by predicted modelling studies and function of the human ortholog [30,31]. From Hsp70 and Hsc70 class of proteins, Hsp70Aa was observed to suppress the mutation whereas Hsc70Cb, and Hsc70-4 enhanced the mutation.

To quantitate the effect of chaperones on the growth of the mutant we monitored a 48 hrs growth curve. Huntington mutant 103Q with empty vector Luciferase and those co-expressing with the chaperones were grown in SD media. The cultures were treated with raffinose for removal of glucose from the media and induced with 2% galactose. The cultures were grown in galactose medium for 48 hrs in a 96-well plate in triplicates. O.D. was measured every 30 minutes with orbital shaking at 30^0^C and the graph was plotted using Graph pad software. In Fig.1B, the blue line indicates the growth curve of HTT103Q with empty vector and the red line denotes growth curve of the mutant in presence of the respective chaperones. The area under the curve of all the chaperones including Luciferase were calculated. They are then converted to a logarithmic scale. The logarithmic values of chaperones were normalized with the logarithmic values of Luciferase. Chaperones which made the mutant grow faster than empty vector Luciferase had more area under the curve in comparison to Luciferase. These were aligned towards the right side of Luciferase with positive values as shown in Fig.1C. They were categorized as suppressors of the mutation. Chaperones that grew slower than empty vector Luciferase showed a reduced area under the curve. These were aligned on the left side of Luciferase after normalization. These showed negative values in Fig.1C and were collectively termed as enhancers of the mutation. The sequence of suppressors and enhancers in order of proximity to Luciferase is given in Table 1.

**Fig1.**
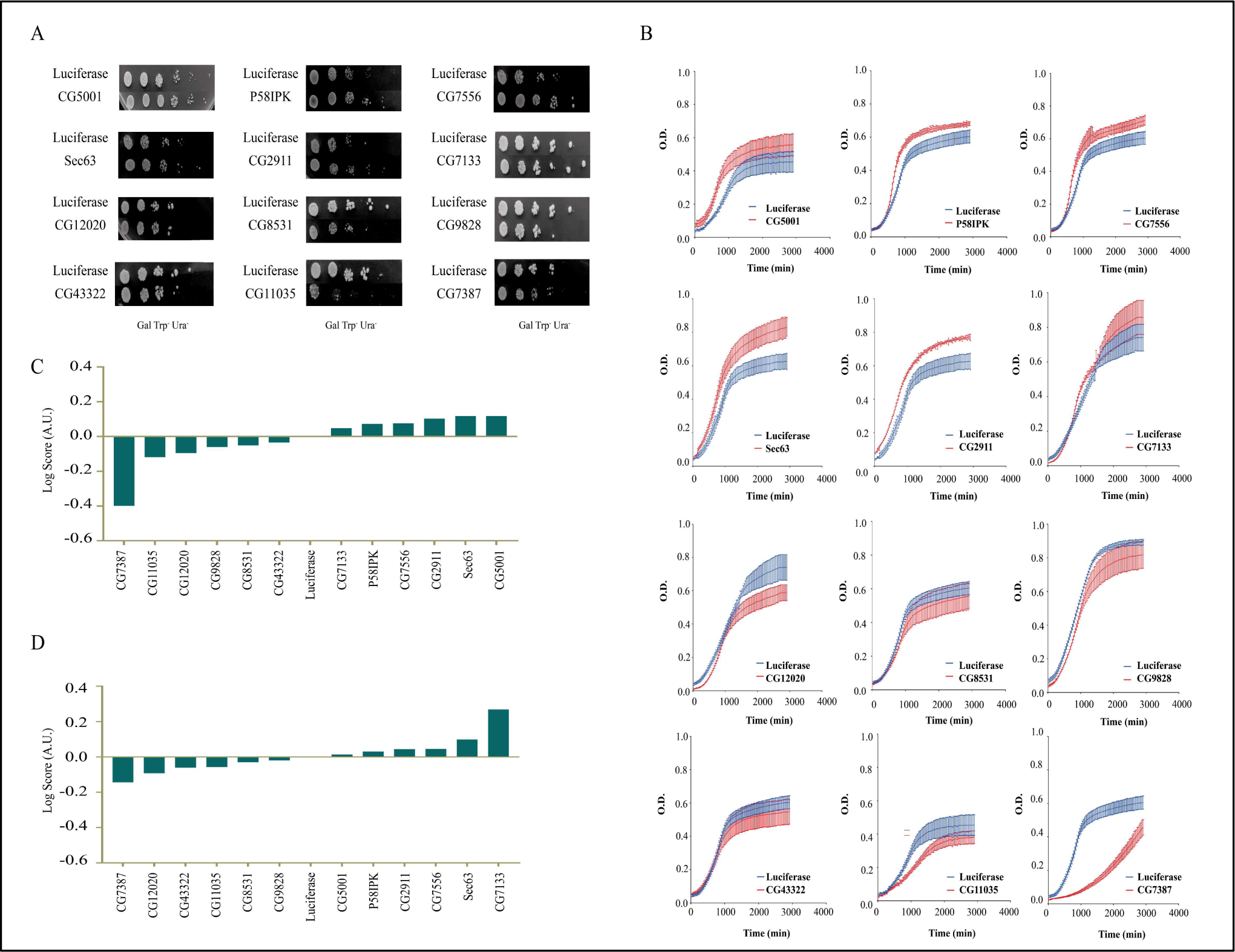
Effect of DnaJ chaperones on growth of Huntington’s mutant. A. DnaJ domain chaperones transformed in *Saccharomyces cerevisiae* model of Huntington’s muant 103Q serially diluted and spotted on Gal Trp^-^Ura^-^ plates. Empty vector Luciferase serves as control. B. 48 hours growth assays of mutant 103Q transformed with DnaJ chaperones (P<0.05). Blue line indicates growth of empty vector Luciferase whereas red line indicates growth curve of respective chaperones. C. Classification of DnaJ chaperones as enhancers and suppressors based on area under the curve values of growth curves, determined using graphpad prism software. Values of area under the curve were converted to a logarithmic scale and were normalized with the slope value of Luciferase. Chaperones to the right of Luciferase are suppressors whereas the ones to the left are enhancers of the mutation. D. Classification of DnaJ chaperones as enhancers and suppressors on the basis of slope of growth curves determined using graphpad prism software. Slope values and area under the curve were converted to logarithmic scale and were normalized with the slope value of Luciferase. Chaperones to the right of Luciferase with positive values are suppressors whereas the ones to the left with negative values are enhancers of the mutation.

**Table 1:**
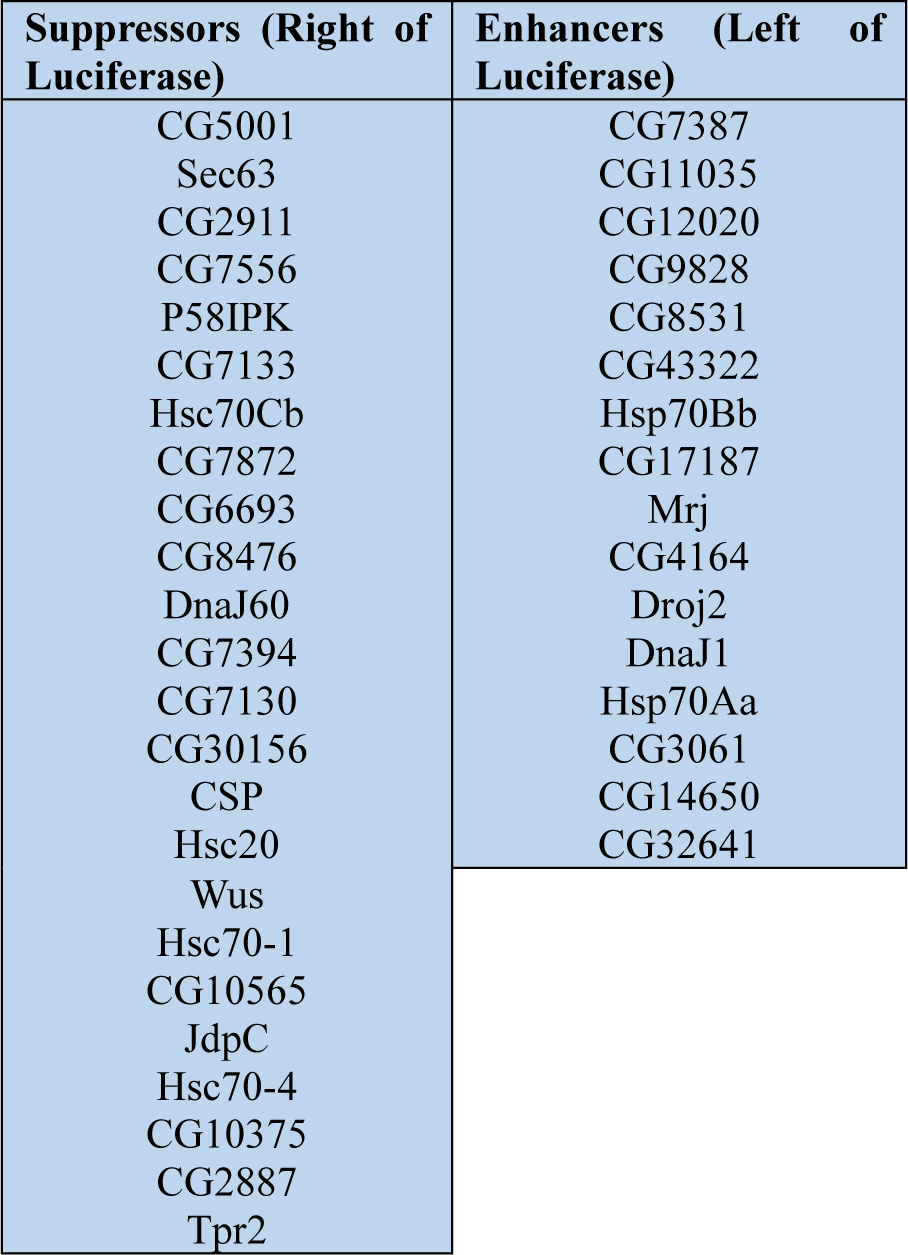
Chaperone classification w.r.t Luciferase for enhancers and suppressors according to area under the curve values.

Similarly, the slope was calculated for all the chaperones to represent the growth rate. The slope values were converted to a logarithmic scale and normalized against the logarithmic slope value of Luciferase. In Fig 1.D suppressors were the ones that had higher slope values than Luciferase and were aligned to the right. Enhancers having reduced slope values than Luciferase were aligned to the left of Luciferase. The sequence of suppressors and enhancers from lowest to highest slope values are depicted in Table 2.

**Table 2:** Chaperone classification w.r.t Luciferase for enhancers and suppressors according to slope value.

| Suppressors (Right of Luciferase) | Enhancers (Left of Luciferase) |
| --- | --- |
| CG7133 | CG7387 |
| Sec63 | CG12020 |
| CG7556 | CG43322 |
| CG2911 | CG11035 |
| P58IPK | CG8531 |
| CG5001 | CG9828 |
| CG7872 | Mrj |
| Hsc70Cb | CG17187 |
| CG6693 | Hsp70Bb |
| CG8476 | CG32641 |
| Hsc20 | Hsc70-4 |
| CG10565 | Hsp70Aa |
| CSP |  |
| CG7130 |  |
| CG4164 |  |
| Wus |  |
| JdpC |  |
| DnaJ60 |  |
| Tpr2 |  |
| CG7394 |  |
| CG30156 |  |
| Droj2 |  |
| CG14650 |  |
| CG3061 |  |
| Hsc70-1 |  |
| DnaJ1 |  |
| CG10375 |  |
| CG2887 |  |

### Chaperones ameliorate protein aggregation in Huntington’s mutant in *S. cerevisiae*

Protein aggregation is a characteristic feature of the trinucleotide repeat in neurodegenerative disorders like Huntington’s. Mutant 103Q transformed with chaperones and empty vector Luciferase were grown in dextrose medium, followed by raffinose wash for removal of residual dextrose and induced by 2% galactose for switching on the htt genes. The cells were then observed for GFP expression under confocal microscope [77]. Wild type 25Q shows GFP diffused all over the cytoplasm whereas mutant 103Q shows punctae of different morphologies, which is an indicator of aggregated proteins. Around 100-150 cells of mutant HTT103Q transformed with each of the chaperones and Luciferase were independently scored. Representative images are shown in Fig.2A. Based on the confocal images, cells were classified into 6 main morphologies, namely, cells with diffused cytoplasm, diffused cytoplasmic punctae, single small punctae, single large punctae, fibre like punctae and multiple punctae (Fig. 2B). Each of these phenotypes relate to a specific toxicity level [78]. Most toxic is the multiple punctae phenotype which have been assigned a toxicity rank of 1. Next is the fibre like punctae phenotype with toxicity rank 2. It is followed by the single large puncta with toxicity rank 3. The single small punctae phenotype is assigned a toxicity rank of 4, diffused cytoplasm with punctae phenotype has toxicity rank 5, and finally the diffused cytoplasmic phenotype is the non toxic form with rank 6. (Fig. 2B) Fig 2B shows the percentage of the various cell types as classified by image analysis, in the Huntington mutant background harboring the different types of chaperones. Weighted mean formula was used to give a specific toxicity score to each of these chaperones [79]. This was categorized by using the number of cells showing the characteristic morphology. The scores of the chaperones were then normalized to Luciferase values. Accordingly, suppressors and enhancers of HD based on chaperone score are as follows:

**Fig2.**
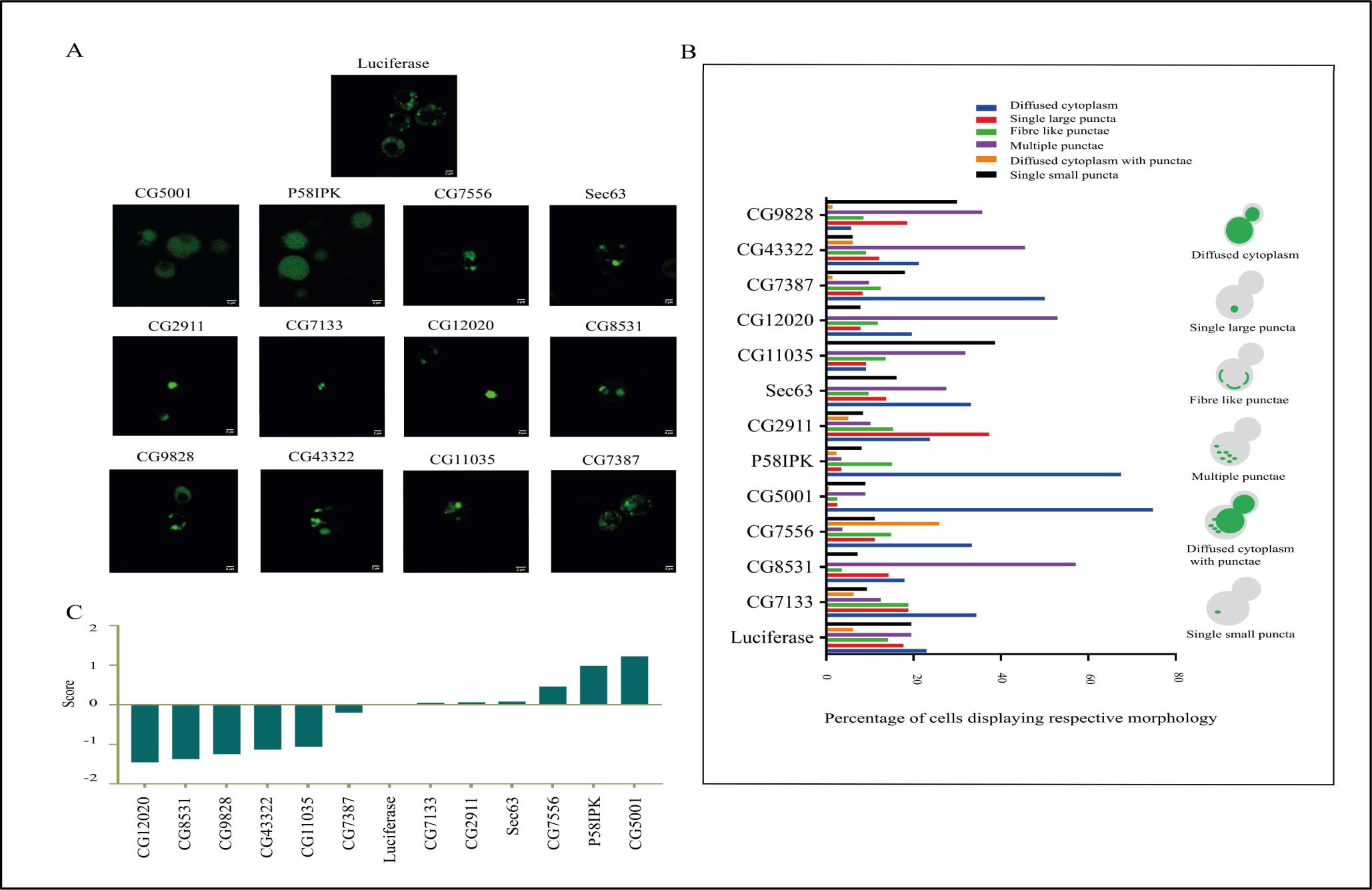
Effect of DnaJ chaperones on protein aggregation of Huntington’s mutant. A. Protein aggregation in *Saccharomyces cerevisiae* model of Huntington’s mutant 103Q transformed with DnaJ chaperones. Aggregates were seen in the form of bright green GFP punctae upon inducing the cells with 2% galactose. B. Different morphologies exhibited by cells with GFP tagged mutant Huntington plasmid 103Q and co-transformed with DnaJ chaperones. C. Classification of DnaJ chaperones suppressing or enhancing Huntington toxicity based on imaging assay. Chaperone score was assigned using weighted mean formula and different toxicity levels to each of the morphologies displayed by the cells. The score of chaperones were normalized against score of Luciferase and the graph was plotted with suppressors having positive score values and enhancers with negative score values.

**Table 3:**
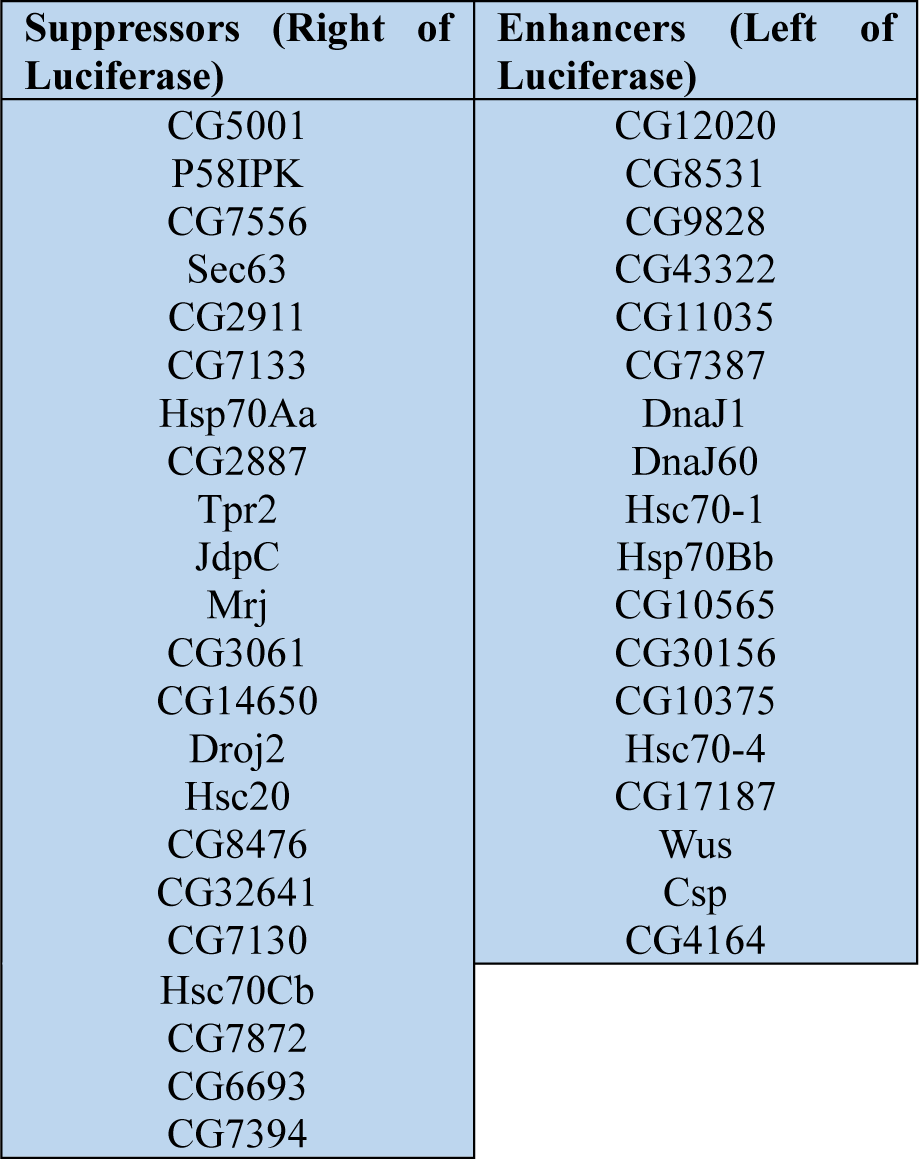
Chaperone classification w.r.t Luciferase for enhancers and suppressors based on microscopy score.

Of all these 40 chaperones, DnaJ60 is an outlier which shows different pattern for growth and imaging assays. It acts as an enhancer in the microscopic data and as a suppressor for the growth curve and serial dilution assay.

### Selection of best fit candidate Enhancers and Suppressors for Huntington’s pathogenicity in yeast

Screening of the best fit enhancers and suppressors was carried out by combining the results from area under the curve and slope values along with the chaperone score assigned via imaging techniques. Accordingly, after the enhancers and suppressors were screened, we arrived at these six potential enhancers and suppressors given below.

**Table 4:** Best fit Enhancers and Suppressors.

| Suppressors | Enhancers |
| --- | --- |
| CG5001 | CG12020 |
| P58IPK | CG8531 |
| CG7556 | CG9828 |
| Sec63 | CG43322 |
| CG2911 | CG11035 |
| CG7133 | CG7387 |

Amongst the suppressors, CG5001 and P58IPK were selected for further analyses as these were most effective in reduction of toxicity of Huntington’s pathogenicity through different analyses.

### CG5001 and P58IPK reduce protein aggregation in yeast

Semi-denaturing detergent agarose gel electrophoresis (SDD AGE) is a technique used to characterize large polymers which usually cannot pass through the pores of acryl amide gels. SDS insoluble aggregates are separated with the help of agarose gel with 0.1% SDS. The proteins are then transferred to a nitrocellulose membrane by capillary transfer. Fig. 3A shows the 25Q construct with a monomeric band present in the SDD AGE. On the contrary, Huntington 103Q mutant shows a polymeric smear. The smear is an indication of protein aggregation in the mutant. In presence of the chaperones CG5001 and P58IPK, smear of 103Q decreases in intensity in comparison to the mutant Huntington as analysed by densitometric scans. This possibly suggests that the insoluble protein aggregation of the mutant reduces in presence of these chaperones. Fig 3B is the corresponding western blot which shows the expression of the Huntington protein with those strains coexpressing CG5001 and P58IPK. Chaperones CG5001 and P58IPK were observed to reduce the toxicity related to Huntington’s mutation by rescuing the slow growth and protein aggregation phenotype. This was verified by spot dilution, growth curve assay, microscopy, SDD AGE, western blot, and separation of soluble and insoluble fractions of the protein. On the basis of these observations, one can possibly draw a relation between the selected chaperones and Huntington aggregates. To further explore if the chaperones have a direct effect on the Huntington aggregates or if it is some indirect pathway that results in the rescue of the Huntington phenotype, a co-immunoprecipitation assay was carried out.

**Fig3.**
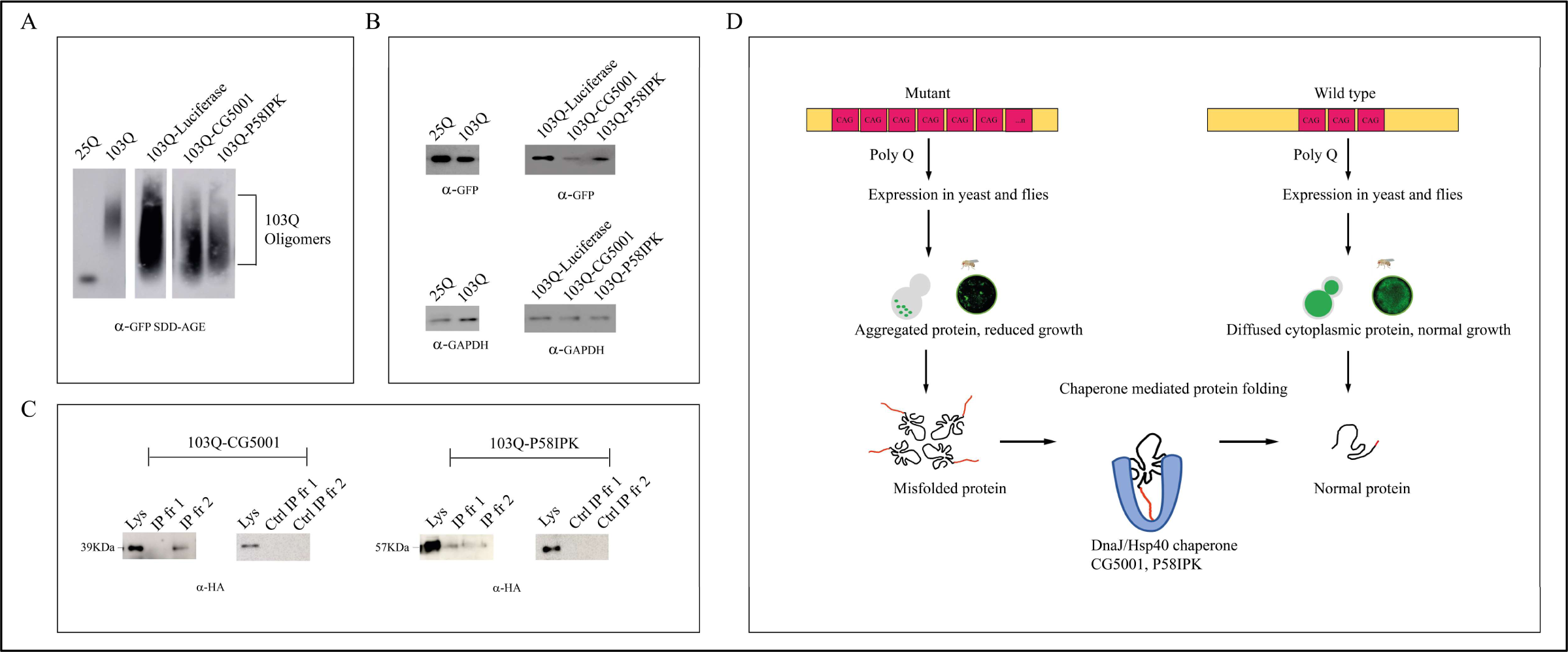
Effect of chaperones CG5001 and P58IPK on protein aggregation using electrophoretic techniques. A. Representative semi-denaturing detergent agarose gel electrophoresis (SDS-PAGE) showing protein aggregation of the mutant. B. Western blot of mutant 103Q with chaperones CG5001 and P58IPK and empty vector Luciferase probed with anti-GFP antibody. C. Representative blot of co-immunoprecipitation of chaperones CG5001 and P58IPK with HTT103Q respectively. D. **Hypothetical Model.** Schematic of the model originating from this work. We hypothesize association of DnaJ domain chaperones with the misfolded proteins that are formed due to repetitive CAG (103Q) repeats, in yeasts and flies, in the pathogenic Huntingtin. The misfolded proteins bind with the chaperones CG5001 and P58IPK causing them to act as an inhibitor of protein aggregation which reduces the toxicity related to multiple CAG repeats.

Co-immunoprecipitation is one of the standard methods for identifying or confirming the occurrence of protein-protein interactions *in vivo*. Huntington constructs used in this study have a GFP tag whereas chaperones proteins are with HA epitope tags. To see the interaction of Huntington aggregates and chaperones, an interaction between 103Q-GFP and respective HA-tagged chaperones CG5001 and P58IPK were checked. Mutant 103Q and 103Q transformed with CG5001 and P58IPK respectively, were grown according to standard conditions, and protein lysates were made by bead beating method. These lysates were incubated with anti-GFP antibody and protein A beads. Proteins were eluted from the beads and run on SDS PAGE gel which was further probed with anti-HA antibody. If chaperones directly interacted with Huntington aggregates, they would be a part of the complex and would be eluted with GFP proteins. It would be possible to detect them on an immunoblot when it is probed with anti-HA antibody. Here, we see bands in the Input and IP lanes of mutant 103Q with CG5001 and P58IPK whereas the negative controls show no bands in the IP. This possibly gives an indication to the respective chaperones CG5001 and P58IPK interacting directly with the Huntington (103Q) protein in yeasts.

### CG5001 and P58IPK facilitate protein conversion from insoluble to soluble fraction in yeast

Huntington aggregates are usually SDS insoluble in nature [80]. A study of Huntington’s disease in mice cortex reported that high molecular weight species of mutant HTT increases significantly in the insoluble fraction of the protein as the disease progresses whereas soluble fractions show a decrease [81]. The soluble and insoluble fractions of HTT103Q and the ones with the chaperones CG5001 and P58IPK were separated by standard techniques from yeast extracts [65,82]. Silver staining of the corresponding soluble and insoluble protein fractions from the SDS PAGE gel was carried out with HTT103Q with luciferase and in presence of chaperones CG5001 and P58IPK. The gel was scanned and densitometric analysis was done using FIJI software. The intensities of 103Q soluble fractions increased in presence of CG5001 and P58IPK in comparison to Luciferase. The intensities of insoluble fractions decreased in presence of CG5001 and P58IPK as compared to Luciferase (Suppl fig. 6). This suggests that the insoluble protein aggregates are being converted to soluble proteins in presence of the chaperones CG5001 and P58IPK (Supplemental Fig.6).

### CG5001 and P58IPK reduce protein aggregation in S2 cells of *Drosophila*

The chaperones CG5001 and P58IPK were checked in *Drosophila* cell lines, namely S2 cells for their effect on Huntington’s mutation. Various cellular and molecular tools have been developed or adapted in fly research. The complete genome sequence of *Drosophila* is reported. The fly life cycle is short and it is easy to grow a comparatively large number of individuals for genetic, molecular, and biochemical analysis. *Drosophila* S2 cell line is easy to grow and maintain in the laboratory, highly susceptible to transfections and gene knockouts using RNAi, and is well suited to high resolution light microscopic assays.

Both CG5001 and P58IPK chaperones from *Drosophila* were selected from a class of DnaJ chaperones as they were rescuing the growth defects and aggregation of the proteins in the screening assay of the yeast Huntington model. Based on these findings, it can be considered that CG5001 and P58IPK are suppressors that help to control the toxicity of the pathogenic aggregates. The human homologs for CG5001 and P58IPK are DNAJB5 and DNAJC3 respectively [82–84]. It is interesting to note here that DnaJB class of chaperones have been reported to play a crucial role in protection against neurodegeneration by maintaining neuronal proteostasis both in *vitro* and *in vivo* [84,85].

The Huntington disease model was constructed using the *Drosophila* embryonal S2 cell line. 103Q Huntington plasmid and chaperones CG5001 and P58IPK were transfected in S2 cells along with the empty vector (So; Sine oculis) using Gal4, DNA and effectene reagent [62]. 103Q has a GFP tag and chaperone proteins have a HA tag at the C terminal. S2 cells expressing 103Q and 103Q with CG5001 and P58IPK and the empty vector were fixed with 4% PFA. Cells were blocked with 5% NGS and probed with anti HA antibody. Immunostaining was carried out using secondary antibody tagged with Alexa fluor 555 (RFP) [28,65]. Upon staining, Huntington aggregates had a GFP tag, and chaperones CG5001 and P58IPK appeared red due to the presence of the HA tag. We observed colocalization of the chaperones CG5001 (r=0.8) and P58IPK (r=1) with the Huntington GFP protein. HTT 103Q without chaperones showed the presence of green fluorescent punctae due to insoluble protein aggregation. These green punctae were found to diminish when HTT103Q was cotransfected with the chaperones CG5001 and P58IPK.

Semi-denaturing detergent agarose gel electrophoresis (SDD AGE) helps to separate SDS insoluble HTT oligomers. S2 cells were lysed, centrifuged, and resuspended in buffer containing SDS [85,86]. Gel was prepared using 1X TAE and 0.1% SDS. Proteins were transferred to a PVDF membrane using capillary transfer. The blot was probed with anti-GFP antibody. S2 cells show a polymeric smear of 103Q. The smear intensity decreases in presence of CG5001 and P58IPK in comparison to 103Q alone as shown in Fig. 4F. (103Q with CG5001 and P58IPK shows a reduction in smear intensity, compared to control) This suggests reduced protein aggregation in the presence of chaperones. Immunoblots reveal, that the Huntington protein is expressed at almost the same level in CG5001 and P58IPK constructs in comparison to the wild type.

**Fig4.**
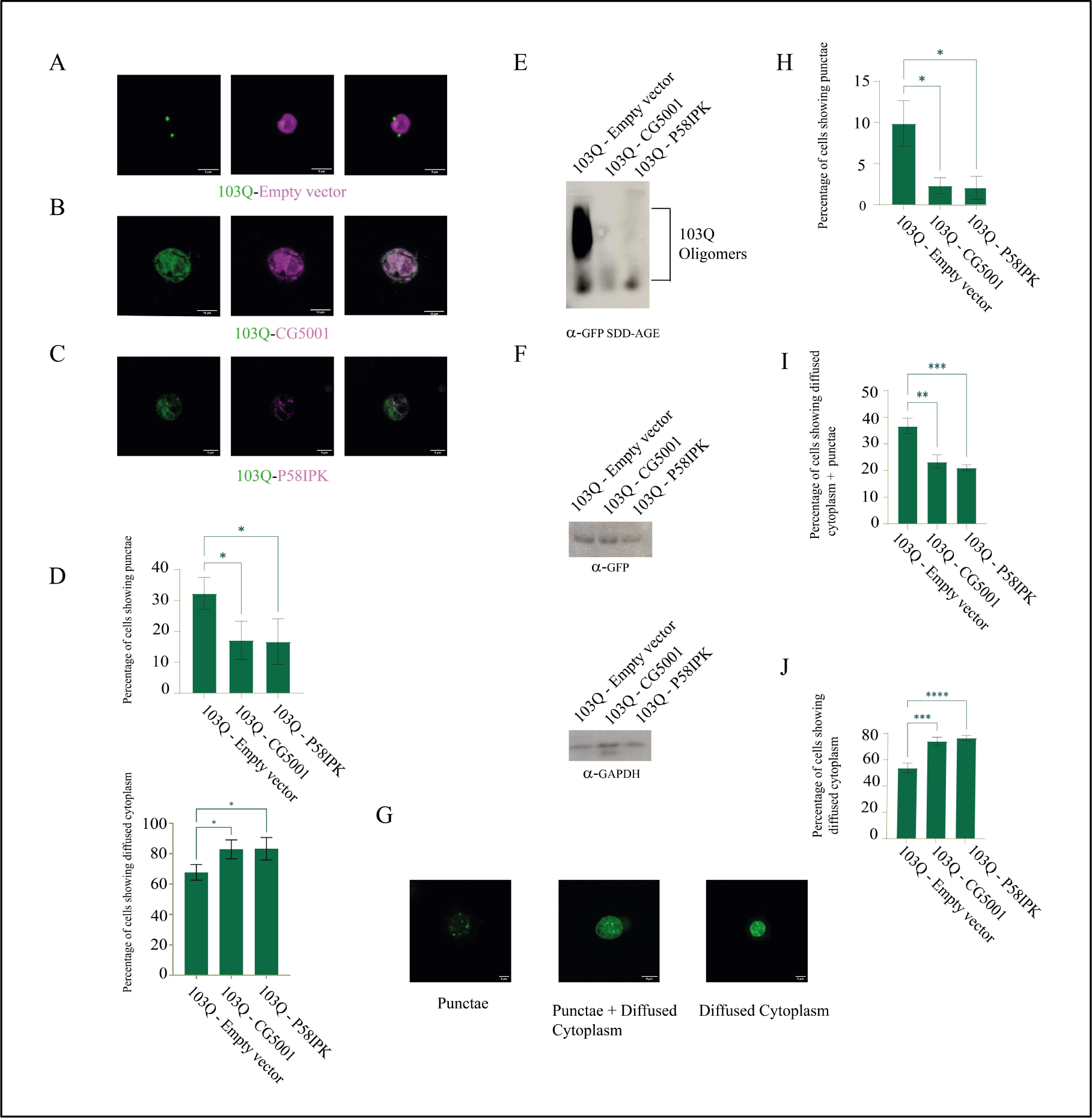
Effect of CG5001 and P58IPK on Huntington mutation 103Q in *Drosophila* S2 cells. A. Transfection of 103Q (green) and with empty vector having HA tag (magenta) in S2 cells. B. Mutant 103Q-S2 cells transfected with CG5001 and immunostained with RFP (shown here in magenta) show colocalization of GFP and RFP. C. Mutant 103Q-S2 cells transfected with P58IPK and immunostained with RFP (shown here in magenta) show colocalization of GFP and RFP. D. Representative graph of S2 cells when transfected with 103Q and chaperones CG5001 and P58IPK respectively shows less protein aggregation punctae as compared to 103Q alone. E. SDD-AGE analysis of mutant 103Q-S2 cells shows decreased protein aggregation in presence of chaperones CG5001 and P58IPK as compared to 103Q when probed with anti-GFP antibody. F. Western blot of mutant 103Q-S2 cells transfected with CG5001 and P58IPK respectively probed with anti-GFP antibody. G. Larval hemocytes from fly crosses of 103Q and chaperones CG5001 and P58IPK with 103Q were observed under fluorescence microscope. Graphs representing percentage of larval hemocytes expressing punctae (H), diffused cytoplasm+punctae (I) and diffused cytoplasm (J) only.

Following the results of the chaperones CG5001 and P58IPK in *Drosophila* S2 cell lines for their effect on Huntington’s mutation, we decided to observe their effects on the aggregation of Huntington’s protein in flies. For this purpose, a transgene *Drosophila* line with 103Q was coexpressed with CG5001 and P58IPK respectively. The Cg Gal4 line was used to express the transgenes in the hemocytes using Gal4-UAS system. Larval hemocytes were dissected and observed under a fluorescence microscope. We found three different morphologies of aggregates in the mutant 103Q and those expressing CG5001 and P58IPK respectively. Upto 7 independent biological replicates were imaged for quantitation for each set of transgene cross. The number of puncta (in both categories, solely puncta and puncta with diffused cytoplasm morphology) was reduced in cell lines with chaperones as compared to the cell lines with empty vector control (AP2 sigma), whereas cell lines with chaperones showed comparatively greater percentage of diffused cytoplasm and less protein aggregation. This indicates a rescue from HD due to P58IPK and CG5001. The scores assigned were evaluated using GraphPad, and the results were found to be statistically significant (p<0.05). Reduction of GFP punctae in S2 cells and larval hemocytes implies a decrease in the pathogenic aggregates.

### CG5001 also rescues growth defects in ALS mutant strains of *S. cerevisiae*

ALS strains of *S. cerevisiae* with TDP-43 and FUS showed slow growth in spot dilution assay compared to empty vector luciferase. When CG5001 was overexpressed in these mutant strains, the slow growth phenotype was rescued. (Supplemental Fig 7). Thus, CG5001 also acts as a rescuer for another neurodegenerative disease pathogenicity as well.

Studies from other groups have shown that overexpression of DnaJB1 and its yeast homolog Sis1 decreases TDP-43 mediated toxicity by affecting the cell growth, shape, and ubiquitin-proteasome system inhibition [62,65]. Overexpression of DnaJB2 improves muscle performance and motor neuron survival in mutant SOD1, carried out in *in vitro* models of ALS [66].

DnaJB5, the human homolog of CG5001 needs to be explored further, to find the mechanism of suppression of toxicity, related to neurodegenerative diseases.

To summarize, our findings from the initial screen in the Huntington mutant in yeast, we can hypothesize that certain DnaJ domain chaperone proteins are effective in reducing the load of aggregated proteins in Huntington’s disease. Out of which, selective ones like CG5001 and P58IPK, with findings from both yeasts and flies show that they are candidates which have a potential in reducing the pathogenicity in Huntington’s disease. Predicted function of CG5001 from modeling studies show that it acts as a chaperone cofactor assisting in protein refolding [30,31] in the cytosol. CG5001 was previously reported to help ataxin-3 suppress Poly Q toxicity in fly eyes caused by Q78 mutation [86,87]. It also has a role in ER stress response. PERK is a Protein kinase R like eIF2α kinase which is normally activated in ER stress response. It works by attenuating protein synthesis and reducing ER client protein load. P58IPK is known to inhibit PERK and plays a functional role in downstream markers of PERK activity in later phase of ER stress response [87,88]. This is specifically interesting as it has been shown that Huntingtin is translationally regulated and the eIF2α translation initiation factor is upregulated in the fly cell lines [54]. Considering these earlier works, we need to check along these lines, to find if other translation associated parameters can be regulated by these two chaperone modifiers in neurodegenerative diseases like Huntington, ALS, Parkinson’s etc.

Multiple sequence alignments of these chaperones reveal that a lot of these chaperones have protein sequences conserved across species of yeasts, *Drosophila*, and humans. (Supplemental Fig.8) This makes these chaperones a potential target for gene therapy for the disease [89].

Hsps are molecular chaperones that are recruited by protein aggregates in human [88,90] and mouse disease models [89–93]. These heat shock proteins have been overexpressed to rescue cell death caused by proteins with abnormal poly Q expansions [92,94]. Hsp104 was found to counteract poly Q toxicity in mammalian neuronal cultures as well as in Huntington’s models of rats and mice [92–96]. HspB8 interacts with the misfolded RNA recognition motif of FUS (fused in sarcoma) and partitions into FUS condensates and prevents their hardening [95,97]. Structural and material properties of HTT fragments of protein in intracellular aggregates can form liquid like reversible assemblies. These assemblies are driven by Huntington’s poly Q tract and proline rich regions. These assemblies are converted to solid fibrillar like structures in cells and *in vitro* [96,98]. It can be hypothesized that CG5001 and P58IPK can emerge as rescuers in similar model studies.

Huntington N terminal fragments have been reported to yield three phases based on aggregate morphologies and sizes [99]. Gaining more insights into the phase separation of Huntington’s aggregates and how these chaperones might influence the aggregates will provide a better understanding of the pathophysiology of HD.

## Conclusion

This study includes yeast and fly cell lines as models of Huntington’s disease and focuses on DnaJ domain chaperones reducing the toxic effects of the mutation. Cloning DnaJ domain class chaperones in yeast mutant of Huntington’s disease leads to either toxicity and protein aggregation or amelioration in the mutant depending on the chaperone. Chaperones that increase the growth rate and decrease the toxic protein aggregation are suppressors of the mutation. CG5001 and P5IPK are the best fit suppressors according to the screen. Huntington mutant S2 cells and *Drosophila* transgenes coexpressing these two chaperones show a substantial decrease in the toxic protein aggregation in the cell.

These chaperones should be explored further for their ability to manage the effects of other neurodegenerative diseases like ALS, Parkinson’s, Spinocerebellar ataxia (SCA), and other toxic Poly Q mutations. Since these chaperones also help to reduce ER stress, they can be used in a dosage sensitive manner to manage ER stress and associated translational defects.

## Materials and Methods

• Library for expression of Hsp40 and Hsp70 co-chaperones in yeast and S2 cells:

*Drosophila* Hsp40 and Hsp70 proteins were cloned by Topo-D-Entr cloning kit (Invitrogen) into Topo-D-Entr vector. This was done post PCR amplification using gene specific primers. Constructs were confirmed by sequencing and were transferred to destination vector pUASg-HA for Hsp40 and pUASt-ccdB-FLAG (pTWF) for Hsp70. Destination vector pUASg-HA had 3X HA tag at C terminal [53,100] and pUASt-ccdB-FLAG (pTWF) had 3X FLAG tag in the C terminal. This was done for S2 cell based assays using LR clonase. For expression of Hsp40 and Hsp70 in yeast cells, Topo-D-Entr clones were cloned in destination vector pAG424-GAL-ccdB-HA with 3X HA tag in the C terminal [33].

- Construction of yeast 25Q, 103Q, TDP-43 and FUS strains: Bacterial Transformation:

DH5α cells were made competent using the calcium chloride method. Huntington plasmid (25Q and 103Q) was added to competent cells. Transformation was done with the help of heat shock method and transformed cells were plated on LB plates containing ampicillin. ALS plasmids pRS426 Gal TDP-43 GFP and 416 Gal-FUS-YFP were also transformed using similar method. These are Addgene plasmids numbered 27476 and 29593 respectively and are deposited by the Gitler lab.

Plasmid Isolation:

Transformed bacterial culture pellet was resuspended in solution I (50mM Glucose, 10mM EDTA and 25mMTris-Hcl). Lysis was done using solution II (1%SDS and 0.2N NaOH). pH was neutralized with solution III (5M Potassium acetate and Glacial acetic acid). Supernatant was subjected to Phenol: Chloroform: Isoamyl alcohol. 100% alcohol was used to precipitate out the DNA.

Yeast Transformation:

*Saccharomyces cerevisiae* cells (w303a, MATa, ade2, his3, leu2, trp1, ura3) were grown in a primary culture of YPD (yeast extract, peptone, and dextrose) media until mid-log phase. Cells were resuspended in 100 mM Lithium Acetate (LiOAc) and divided into aliquots which were treated with 50% Polyethylene Glycol, 1M Lithium Acetate, ssDNA, and plasmid at 42^0^C [101]. The plasmids used were Huntington’s 25Q, 103Q, TDP-43 and FUS-GFP, with the Ura^-^ marker. Cells were plated on SD Ura^-^ media (0.67% yeast nitrogen base, 2%Dextrose, 2% agar, and ∼0.07%Amino acid mixture lacking Uracil) [99,102]. The media are from Clontech Laboratories, Mountainview, CA.

Double transformations were carried out in these mutant backgrounds with different DnaJ chaperone plasmids with a Trp^-^ marker. The final plating was done in SD Trp^-^ Ura^-^ media (0.67% yeast nitrogen base, 2% Dextrose, 2% agar and ∼0.07% Amino acid mixture lacking Uracil and Tryptophan) [100,101]. For every double transformant, 2 representative clones were selected for further assays.

• Serial dilution assay:

25Q and 103Q strains were grown in SD Ura^-^ medium. They were then washed and grown in Raf Ura^-^ medium (0.67% yeast nitrogen base, 2% Raffinose (Raf) and ∼0.07% amino acid mixture lacking Uracil). Optical Density (O.D.) was adjusted to 0.2 and the culture was serially diluted and spotted on SD Ura^-^ and Gal Ura^-^ plates (0.67% yeast nitrogen base, 2% galactose, 2% agar, and ∼0.07% amino acid mixture lacking Uracil) [101,103]. Pictures were taken 24 hours post spot dilution assay on galactose medium.

At least 2 clones from each of these double transformants of 103QHTT with the chaperones were spotted onto SD Trp^-^ Ura^-^ and Gal Trp^-^ Ura^-^ medium in presence of 103QHTT with the empty vector luciferase.

• Growth Assay:

Double transformants of HTT103Q yeasts with and without chaperones were grown in SD Trp^-^ Ura^-^. This was carried out in 3 independent biological replicates with 2 different transformants of Huntington mutant carrying a copy of the different chaperones. Cells were subjected to 2% Raf Trp^-^ Ura^-^ wash and were grown till they reach mid-log phase. 2% galactose was added to the media and cultures were grown in a 96 well plate in triplicates [104]. O.D. was recorded every 30 min. for 48 hours at 30^0^C with continuous shaking in the Epoch2 BioTek Spectrophotometer [102,104]. The data was acquired and growth curve was plotted using graph pad software.

• Microscopy:

25Q and 103Q Huntington mutant yeast strains were grown overnight in S.D. Ura^-^ medium. O.D. was adjusted to midlog phase in Raf Ura^-^ and induced with 2% galactose. After 24 hours, cells were observed under Nikon Ti Eclipse microscope with GFP laser using 60x and 100x objectives. All the images were acquired at 5X zoom using 100X plan Apo objective with 1.49 NA, using the manufacturer’s software. Images were processed using FIJI software [101,105]. For the HTT103Q strain coexpressing the chaperones, cells were grown in SD Trp^-^ Ura^-^ and Raf Trp^-^ Ura^-^ followed by induction with galactose and used for further processing. Number of cells showing variable toxicity were calculated from 3 different biological replicates with cell counts ranging from 100-120 in each category.

Quantitation of Image Analysis:

Mutant strain transformed with different chaperones after image acquisition, shows cells displaying morphologies of diffused cytoplasm (toxicity rank 6), diffused cytoplasm with punctae (toxicity rank 5), single small punctae (toxicity rank 4), single large puncta (toxicity rank 3), fibre like punctae (toxicity rank 2) and multiple punctae (toxicity rank 1) [78] were counted from 3 different representative images of 3 sets of biological replicates for each strain. A score was assigned to each chaperone using the following weighted mean formula.

(% of cells showing 1^st^ morphology ÷ 100) × rank of 1^st^ morphology + (% of cells showing 2^nd^ morphology ÷ 100) × rank of 2^nd^ morphology +…..+ (% of cells showing 6^th^ morphology ÷ 100) × rank of 6^th^ morphology.

The score was then normalized against score of empty vector Luciferase and the graph was plotted.

• SDD-AGE (Semi Denaturing Detergent Agarose Gel Electrophoresis):

Galactose induced yeast cultures were grown till mid log phase. Yeast spheroplasts were made using spheroplasting solution (3M D-Sorbitol, 1M MgCl_2_, 1M Tris, β-mercaptoethanol, Zymolyase) Lysis was done using lysis buffer (1M tris, β-mercaptoethanol, Benzonase, triton-X, PMSF). Protein was quantified using Bradford’s reagent. Gel was prepared using 1.5% agarose in 1x TAE and 0.1% SDS [99,102]. Samples were loaded and run at 55 volts for around 3 hrs. PVDF membrane transfer was set up overnight [103,106]. Blot was developed using ECL substrate (Thermo Fisher Scientific) and developed using Amersham Imager 600 (GE Healthcare Life Sciences). For SDD AGE of S2 cells, they were lysed using SDS and S2 lysis buffer (50mM Tris Hcl, 150mM Nacl, 1% Nonidet P-40), sonicated and then loaded in the gel.

• Western Blot:

Galactose induced yeast cultures were pelleted down after 24 hours of induction. Cell pellet was lysed via bead beating (mention type of bead here with company) using lysis buffer (50mM Tris, 150mM Nacl, 0.1% Nonidet P-40, 1mM DTT, 10% glycerol and protease inhibitor cocktail Roche). The protein was equalized using Bradford reagent and separated using 12% SDS PAGE gel at 120V for around 1.5 hrs. The gel was transferred to PVDF membrane overnight using capillary transfer. Blot was developed using ECL (Thermo Fisher Scientific) and Amersham Imager 600 (GE Healthcare Life Sciences) [57]. For Western blot of S2 cells, they were lysed using SDS and S2 lysis buffer (50mM Tris Hcl, 150mM Nacl, 1% Nonidet P-40), sonicated and then loaded in the gel.

• S2 transfection:

S2 cells were grown in Schneider’s media supplemented with 10% FBS. Transfections were done using effectene and gal4 using manufacturer’s protocol [104,107]. Immunostaining: Transfected S2 cells were added to concanavalin A coated dish and allowed to adhere for 10 min. Excess media was removed and cells were fixed with 4% PFA. PBST washes were given to the cells. Cells were blocked with 5% NGS for an hour. Primary antibody anti-HA was added and cells were kept overnight. They were washed with PBST thrice, and secondary antibody Alexa fluor 555 (RFP) with HA tag was added. Cells were washed again with PBST and were imaged under a Nikon A1 confocal microscope using 60x and 100x objective. In case of chaperones, the 103Q was GFP tagged whereas the antibody was HA specific, which was later stained with RFP. The cells showing GFP and RFP colocalization were observed and quantitated [28,66]. GraphPad Prism (9) was used to analyze and plot the data.

Fly Crosses (screening chaperone lines against Huntington’s disease in *Drosophila melanogaster)*:

The Gal4/UAS system was used to drive gene expression with the following lines: Cg-Gal4 (BDSC stock no. 7011) and UAS-HttQ103-GFP (a kind gift from Prof. Norbert Perrimon’s lab). The Cg Gal4 was used to express the transgenes in the hemocytes. Virgin females of Cg-Gal4 were crossed with males of the UAS-HttQ103-GFP line. The GFP positive virgin females from the progeny were next crossed with the males of two test chaperone lines, UAS CG-5001 and UAS P58IPK and a control, UAS-AP2 sigma (HA tag). Hemocyte dissection was carried out in Schneider 2 (S2) cell media in the GFP-positive larvae of each of these crosses and observed under the confocal microscope. For fixing, larvae were dissected in Schneider 2(S2) cell media, plated on a dish and kept aside for 30 mins for the cells to adhere. The media was then removed and 4% PFA was added and kept for 15 mins. The PFA was then pipetted out and the dishes were washed and stored in 4℃ with 1X PBST. A heterogeneous population was found in the cells. The fields were further quantitated into three categories: cells showing distinct puncta, puncta in a pool of diffused cytoplasm, and cells showing diffused cytoplasm without any puncta. For each trial, around 60-100 cells were observed for each cross.

• Co-immunoprecipitation:

Protein A beads were blocked with TE and BSA overnight at 4^0^C. Anti-GFP purified antibody (Elabscience) was added to the beads (1:1000) and it was kept for binding overnight at 4^0^C. Yeast cell lysate was added to the mixture of beads and antibody for binding overnight at 4^0^C. Beads were washed thrice with yeast lysis buffer. Elution was done by the addition of 0.2 M Glycine (pH 2) and further neutralization with 1.5 M Tris (pH 8.8). The eluates were mixed with loading dye and processed for western blots [54,57]. A mixture of blocked beads and lysate without the antibody was used as a negative control. The membrane was blocked and probed with anti-HA antibody (Proteintech) overnight at 4^0^C and developed using Amersham Imager 600 (GE Healthcare Life Sciences).

• Separation of soluble and insoluble yeast HTT103Q fractions:

HTT103Q yeast cells were grown till mid log phase for 24 hours post galactose induction. Yeast cell pellets were washed with aggregate lysis buffer (ALB). It consists of 50mM potassium phosphate, 1mM EDTA, 5% glycerol, 1mM PMSF, and complete mini protease inhibitor cocktail (Roche). Lysates were then incubated with ALB and 1mg/ml zymolase (Invitrogen). Lysates were then sonicated, centrifuged and supernatant was separated as soluble fraction. Pellet was resuspended in ALB containing NP40 and subjected to sonication. Pellet was washed with ALB and resuspended by sonication. It was then centrifuged and supernatant was stored as the insoluble fraction [108].

• Silver staining:

Soluble and insoluble fractions were run on 12% SDS gel at 120V. Gel was washed in water and fixed using fixing solution (50% methanol, 12% acetic acid, 0.05% formaldehyde) for 1 hour. The gel was washed thrice with 50% ethanol and then washed in 0.8mM sodium thiosulphate for 2 minutes for sensitization with Sodium thiosulfate. Sodium thiosulfate was removed by a water wash and the gel was kept for staining in silver nitrate solution (2mg/ml AgNO_3_, 0.08% formaldehyde) for 15 mins in the dark. It was then washed with water and developed using developing solution (6% w/v Na_2_CO_3,_ 2% 0.8mM sodium thiosulfate, and 0.01% formaldehyde). The gel was put in stop solution (50% methanol, 12% acetic acid) for storage [109].

## Supporting Information

Fig1. Effect of Huntington’s mutation on growth and protein aggregation in yeast cells

Fig2. Effect of chaperones on the growth of Huntington’s mutant studied with the help of spot dilution assay.

Fig3A. Effect of chaperones on the growth of Huntington’s mutant studied with the help of growth curve assay. B. Classification of enhancers and suppressors based on area under the curve. C. Classification of enhancers and suppressors based on the slope.

Fig4. Effect of chaperones on the protein aggregation of the Huntington’s mutant.

Fig5. Serial dilution assay SD control plates

Fig6. Effect of chaperones on soluble-insoluble fractions of Huntington’s mutant

Fig7. Effect of chaperones on growth of yeast overexpressing TDP-43 and FUS as models of Amyotrophic Lateral Sclerosis (ALS).

Fig8. Phylogenetic tree constructed using multiple Sequence alignment of protein coding sequences of chaperones across yeast, humans, and fly.

Fig9. SDD-AGE ratios of intensities of chaperones with respect to empty vector control.

Table1. Hsp40 and co-chaperone gene location and primer set for Topo-D-Entr cloning.

## Acknowledgements

We would like to thank Dr. Renu Mohan for helpful suggestions on preparing the MS. We would like to thank Prof. Michael Sherman for the 25Q and 103Q plasmids and Dr. Jennifer Gerton for sharing yeast strains. The experiments were designed by TB and AM. The yeast experiments were performed by AD, TB, and SA. The fly experiments were performed by RG and AM. MS was prepared by TB, AM, AD, and RG. This work was supported by a research grant from the Department of Biotechnology (DBT-IYBA) (BT/09/IYBA/2015/03), (DBT RLS) (BT/RLF/Re-entry/54/2013) to TB and Wellcome Trust India Alliance Intermediate fellowship to AM (IA/I/13/2/501030). AD and SA are recipients of the ICMR senior research fellowship (ITR-F/2020-ITR).

## Competing Interests

The authors declare no competing interest.

---

**Supplemental Fig1.**
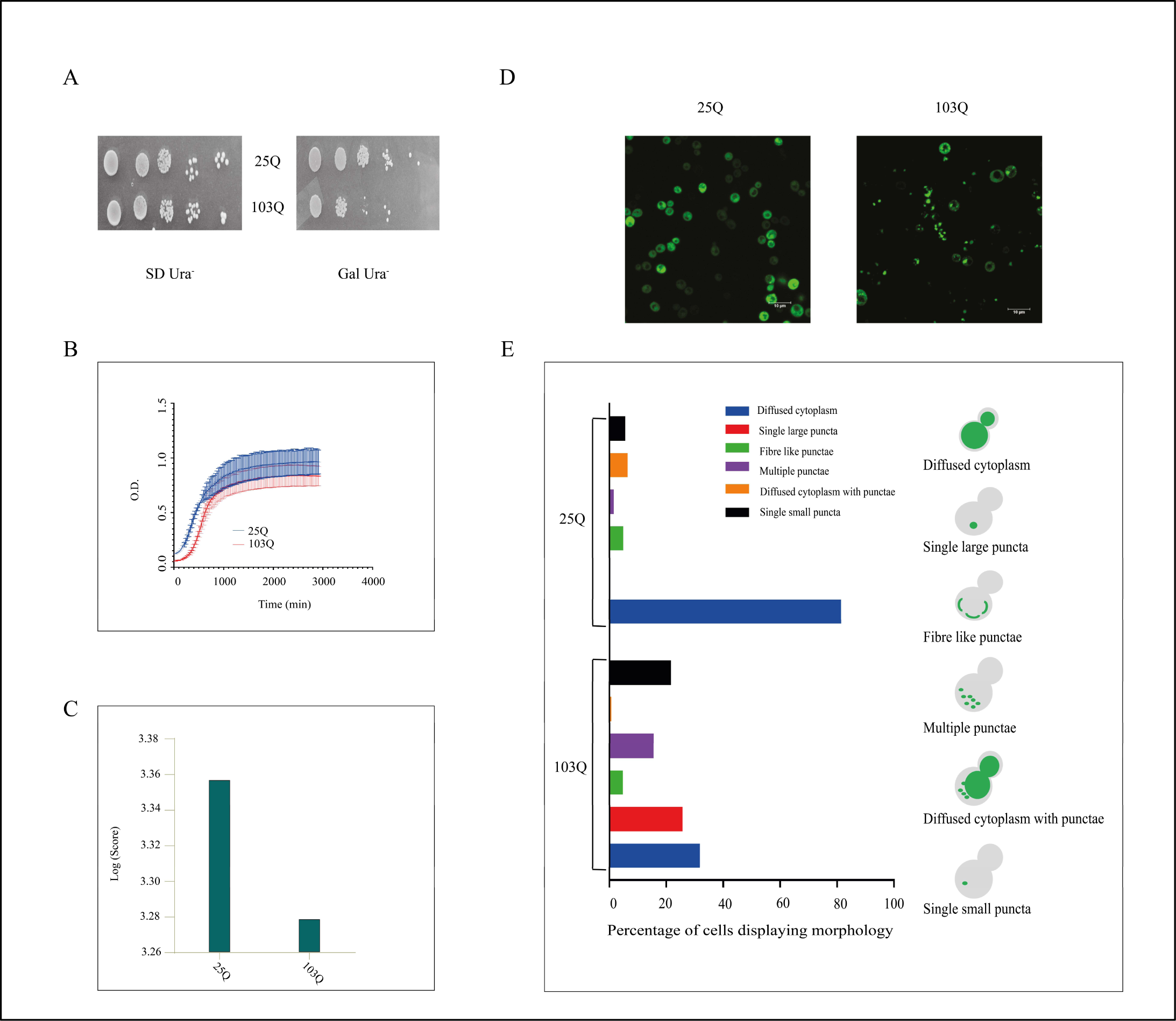
Effect of Huntington’s disease mutation on the growth and protein aggregation of yeast cells: A. Serial dilution assay after transformation of wild type 25Q and mutant 103Q HTT in the host strain *Saccharomyces cerevisiae*. B. 48 hours growth curve assay of 25Q and 103Q by obtaining O.D. every 30 min at 600nm. C. The area under the curve assay for 25Q and 103Q growth curves was converted to a logarithmic scale and plotted. D. Microscopy of 25Q and 103Q GFP under a confocal microscope. E. Analysis of 25Q and 103Q microscopy images categorizing them into different morphologies based on size and nature of protein aggregates. Around 100 cells were counted each for 25Q and 103Q.

**Supplemental Fig2.**
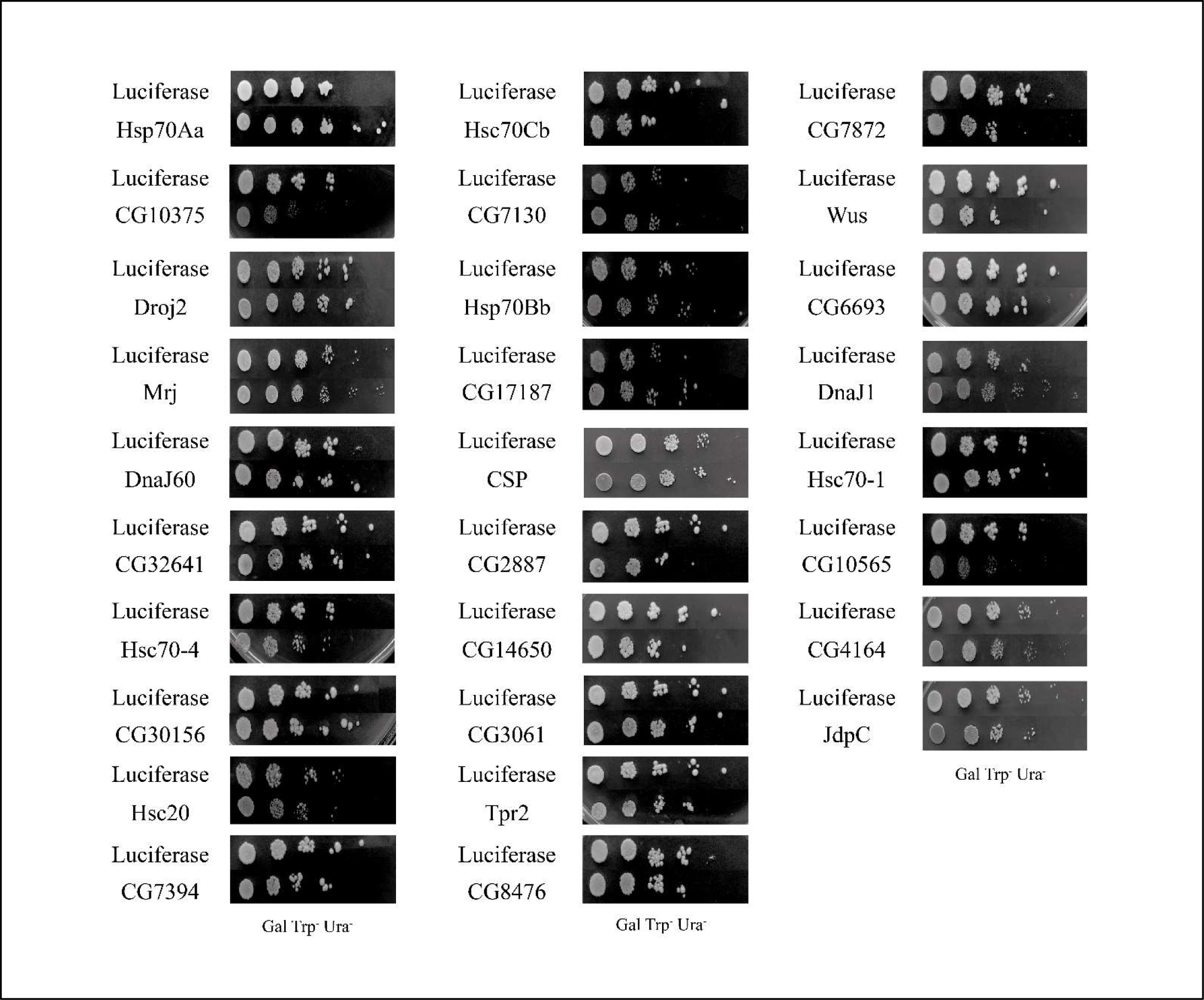
**Serial dilution assay** on Gal Trp^-^Ura^-^ of remaining Hsp40 chaperones described in the study. (Other chaperones shown in Fig.1A).

**Supplemental Fig3.A.**
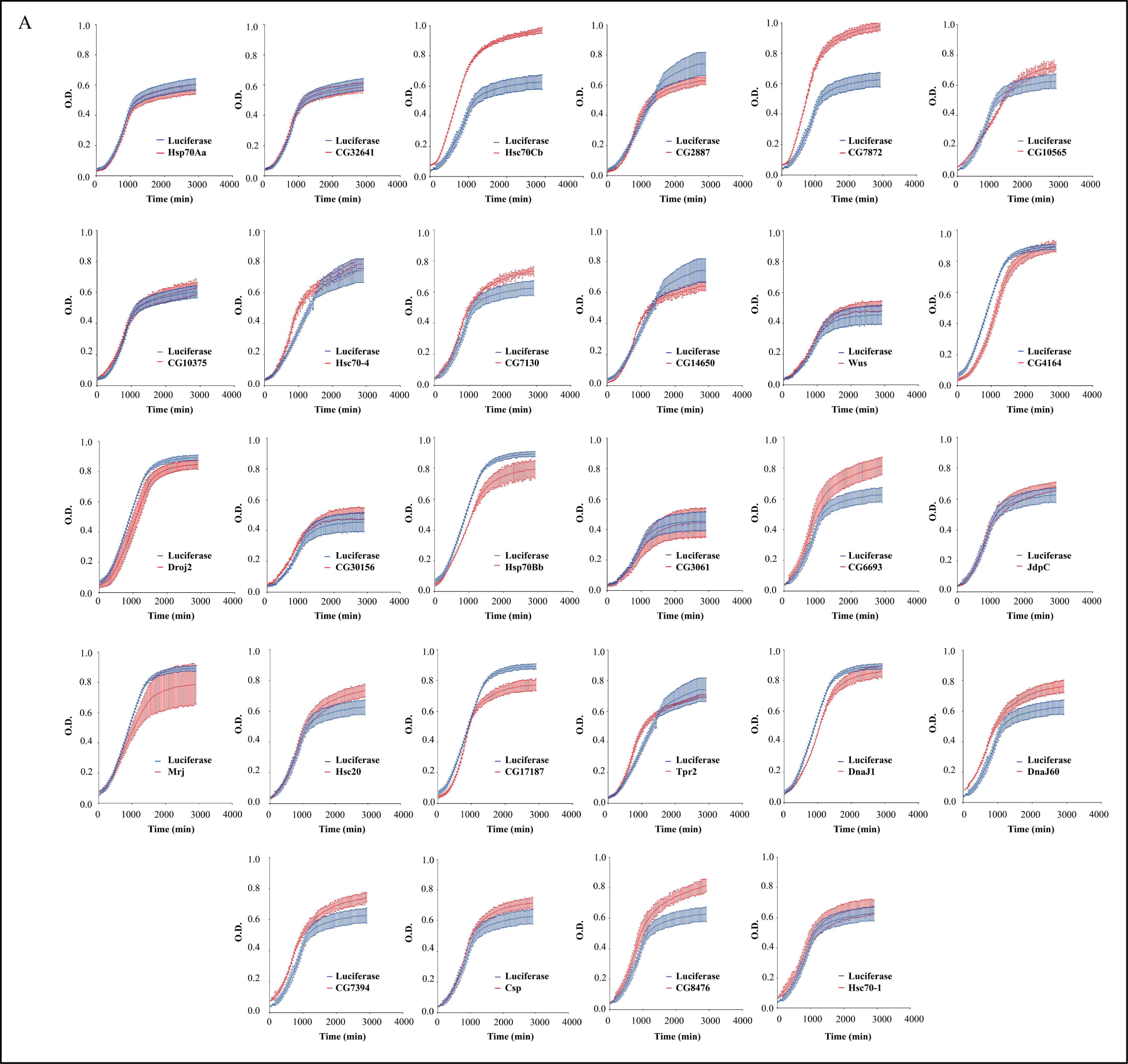
**48 hours growth curve assay** of remaining 28 DnaJ chaperones with galactose induction, discussed in the study. (Others shown in Fig. 1B).

**Supplemental Fig3.B.C.**
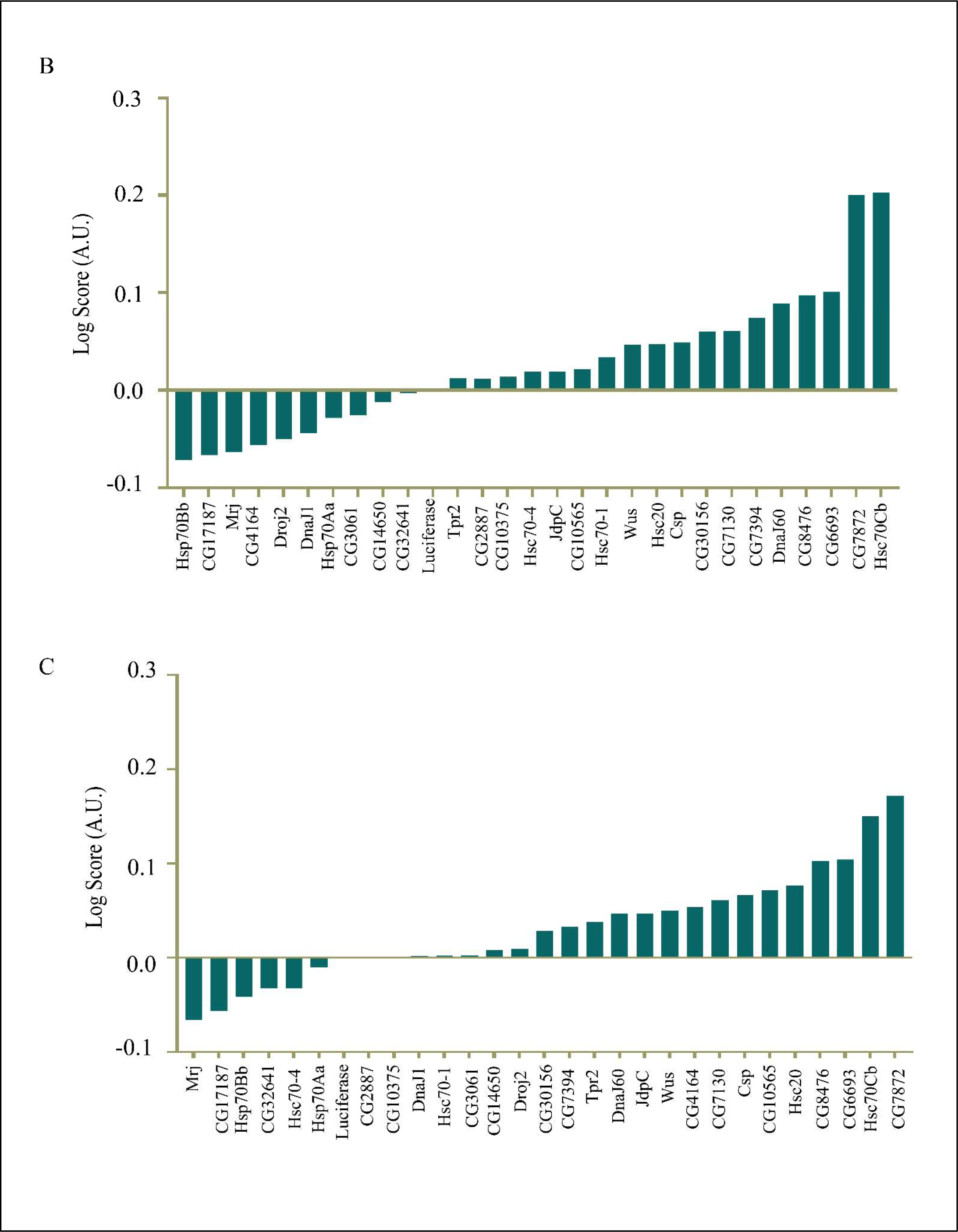
**Classification of DnaJ chaperones** of these chaperones as enhancers and suppressors based on the area under the curve values of growth curves, determined using graphpad prism software. Values of area under the curve were converted to a logarithmic scale and were normalized with the slope value of Luciferase. Chaperones to the right of Luciferase are suppressors whereas the ones to the left are enhancers of the mutation. **Supplemental Fig3.C. Classification of DnaJ chaperones** of the 28 chaperones as enhancers and suppressors on the basis of slope of growth curves determined using graphpad prism software. Slope values and area under the curve were converted to logarithmic scale and were normalized with the slope value of Luciferase. Chaperones to the right of Luciferase with positive values are suppressors whereas the ones to the left with negative values are enhancers of the mutation.

**Supplemental Fig4.**
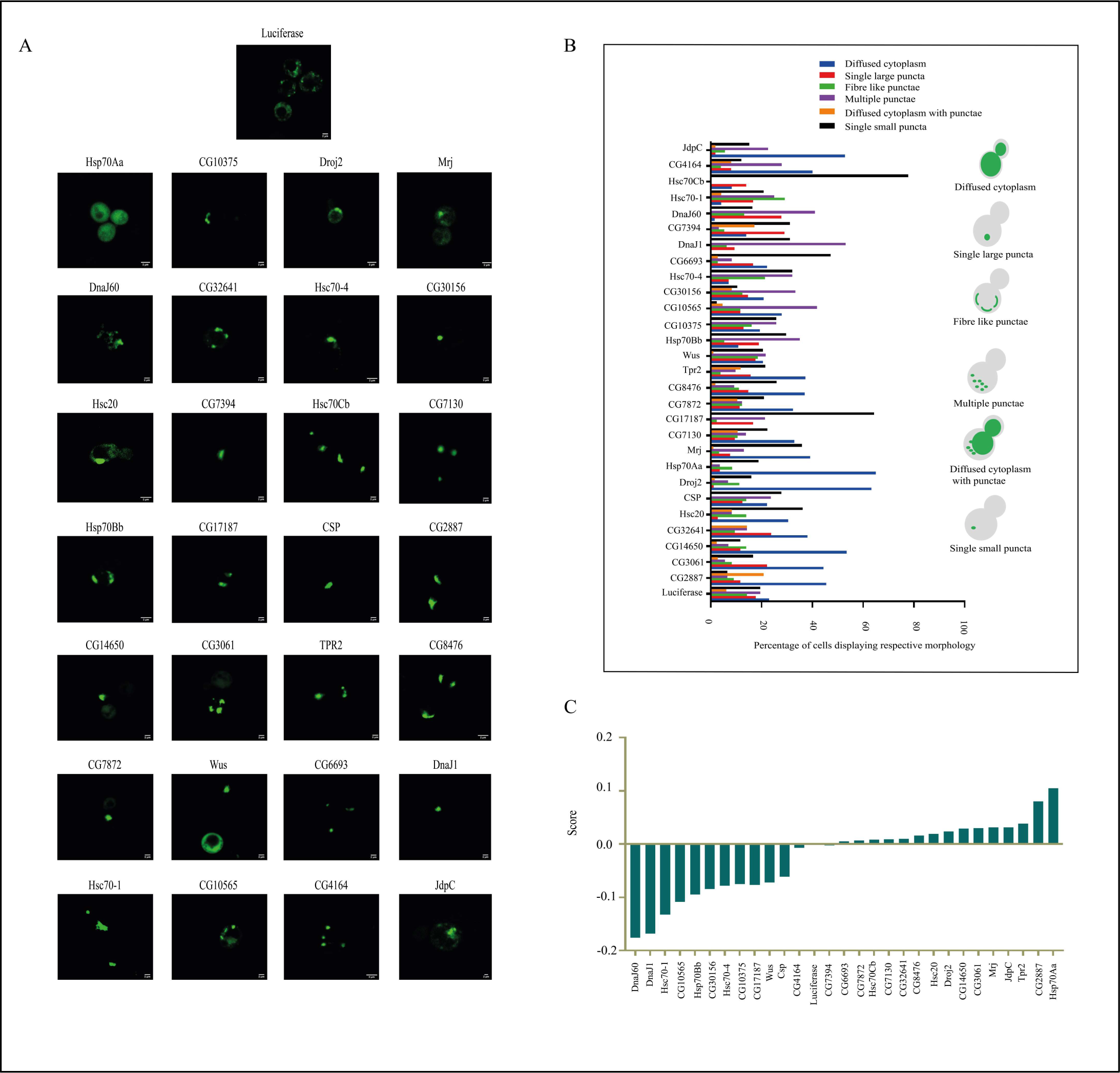
**Effect of 28 DnaJ chaperones on protein aggregation** of Huntington’s mutant. Aggregates are seen as bright green GFP punctae on inducing the cells with galactose (2%). Rest shown in Fig. 2.

**Supplemental Fig5.**
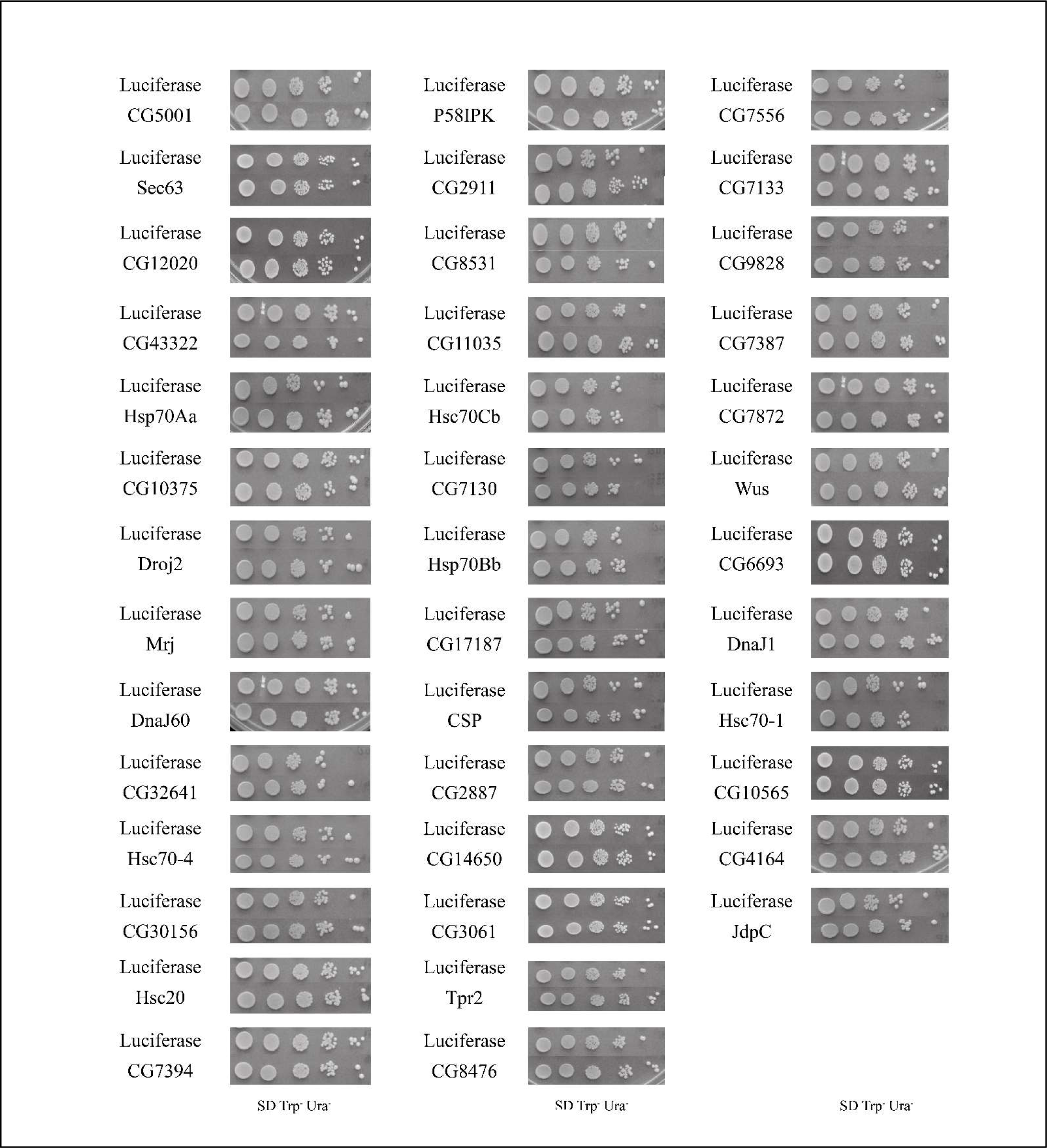
**Effect of chaperones on the growth** of Huntington’s disease mutant in yeast cells by serial dilution assay: Serial dilution assay of Huntington’s disease mutant 103Q transformed with DnaJ domain chaperones and co-chaperones grown on SD Trp^-^ Ura^-^ plates observed after 48 hrs. SD Trp^-^ Ura^-^ plates were used as control as plasmids were galactose inducible.

**Supplemental Fig6.**
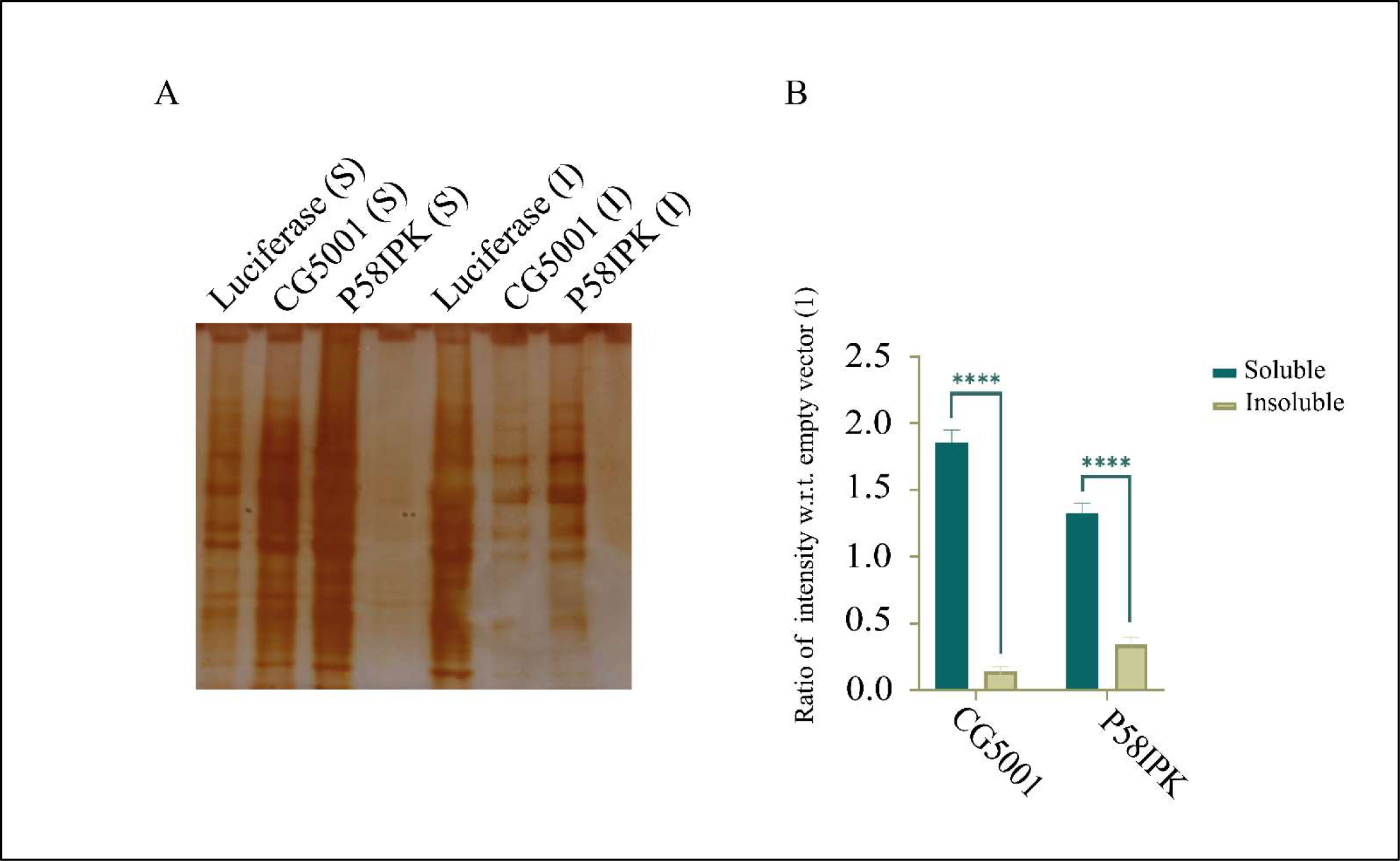
**A. Separation of soluble and insoluble fractions** of mutant 103Q with empty vector Luciferase and chaperones CG5001 and P58IPK was followed by analysis by silver staining. B. Ratio of intensity of soluble to insoluble fractions of CG5001 and P58IPK with respect to Luciferase have been calculated.

**Supplemental Fig7.**
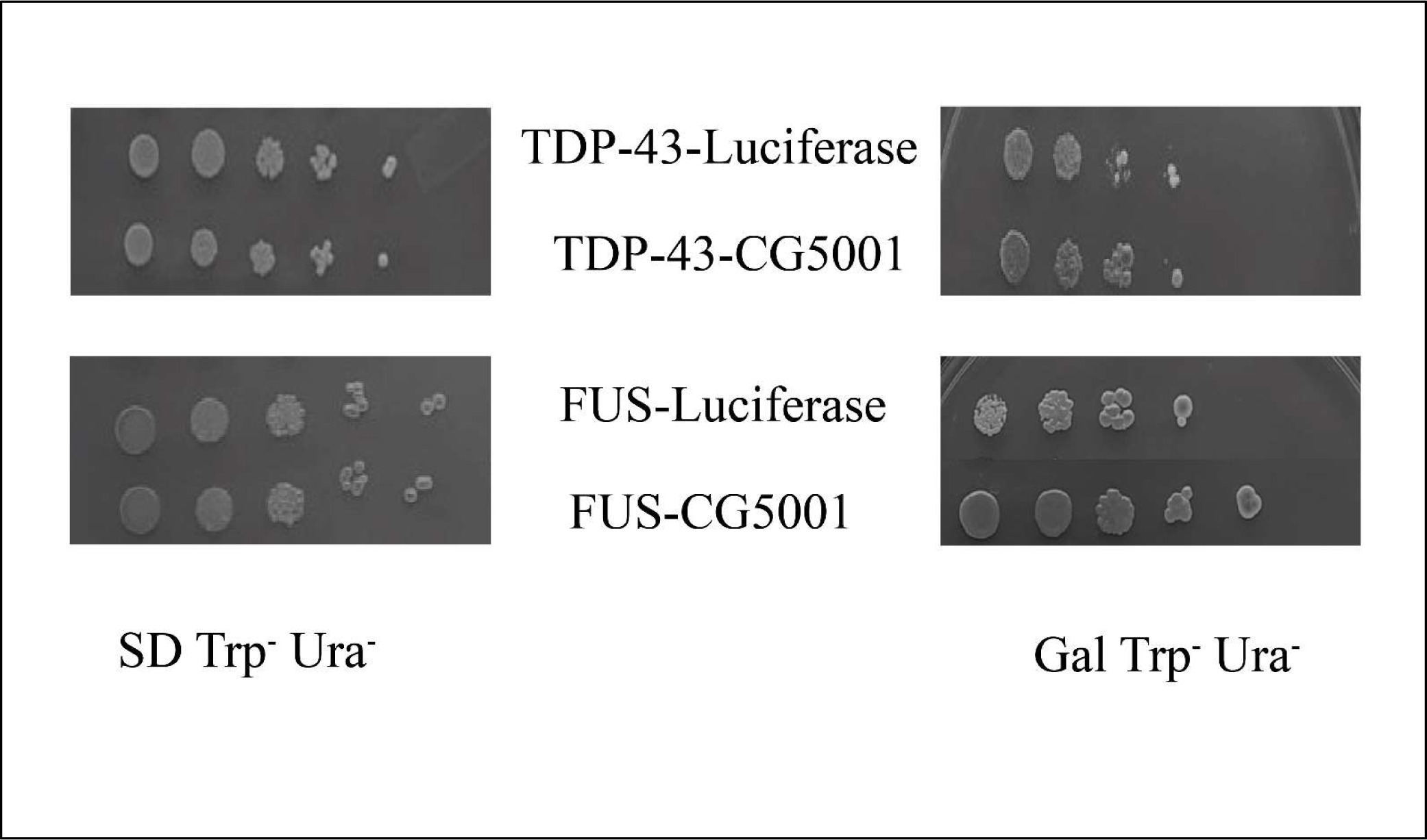
**Effect of chaperone suppressor** CG5001on *S.cerevisiae* strains of ALS mutants TDP-43 and FUS by spot dilution assay on both dextrose and galactose plates.

**Supplemental Fig8.**
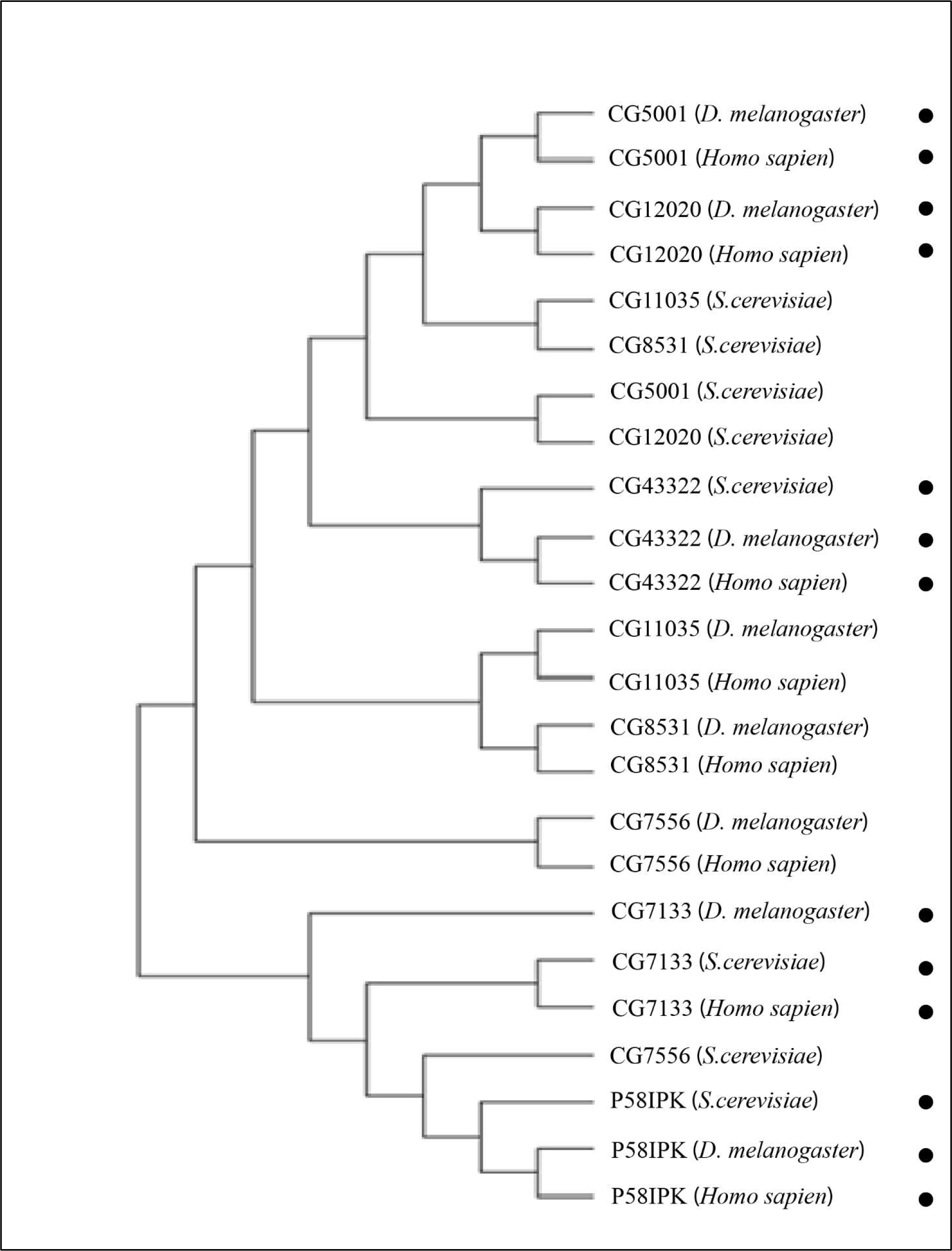
**Phylogenetic tree** constructed using multiple sequence alignment of proteins of chaperones across species with the help of MEGA 11 tool. Hsp40 chaperones showing conserved protein regions across species of yeasts, humans, and *Drosophila*.

**Supplemental Fig9.**
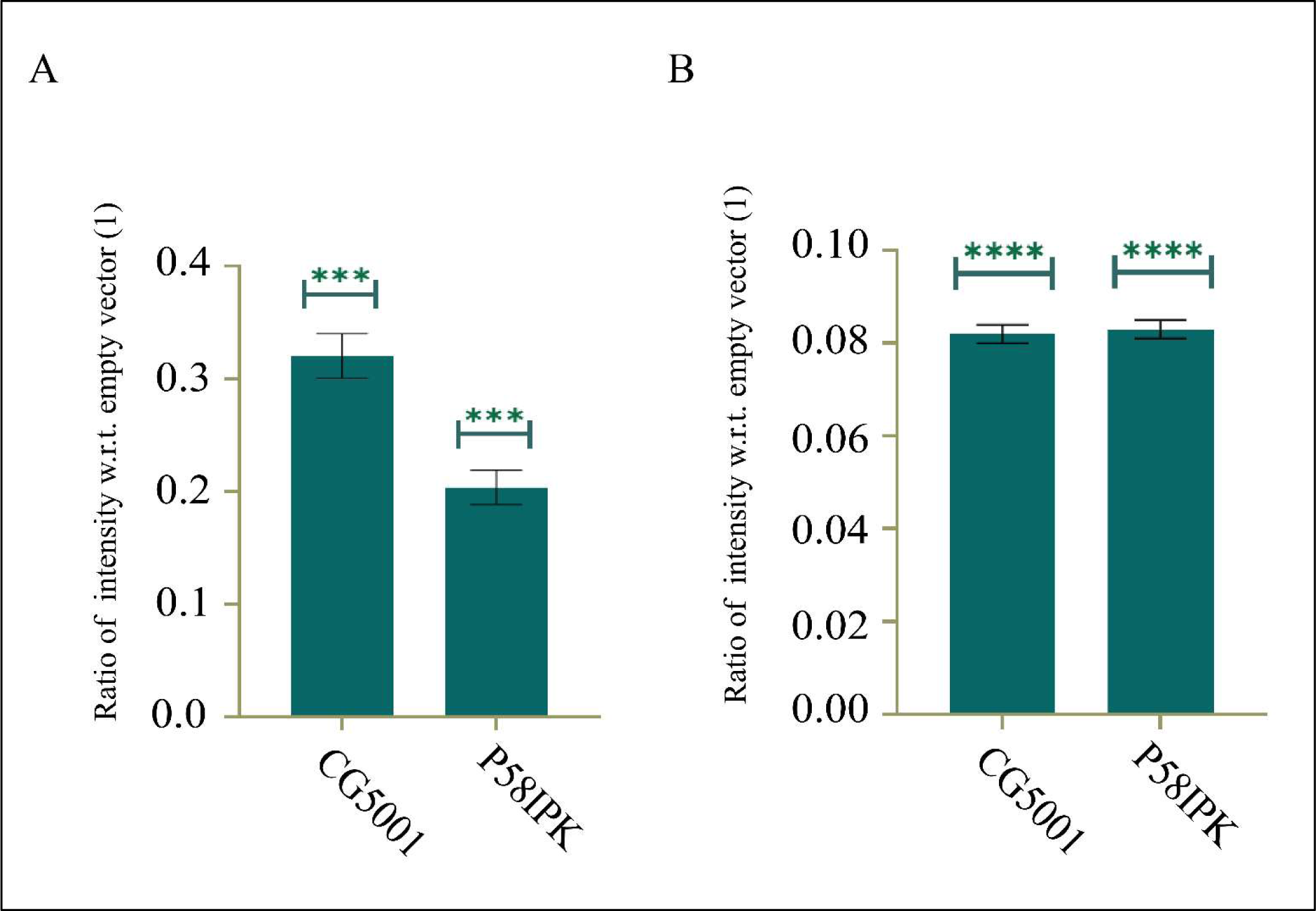
**SDD-AGE Ratios** of intensities of chaperones with respect to empty vector control of *S. cerevisiae* (A) and S2 cells (B).

**Supplemental Table1.**
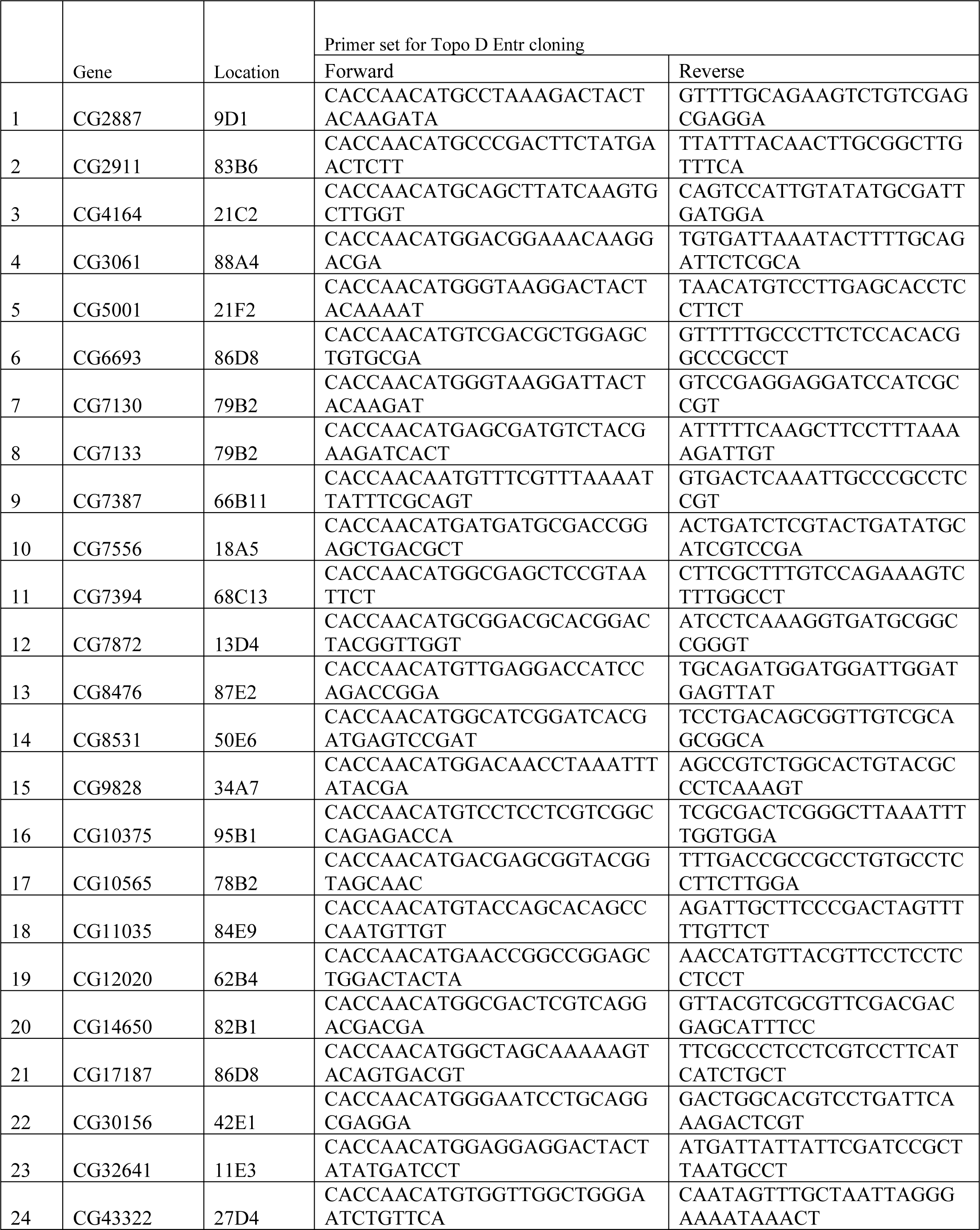
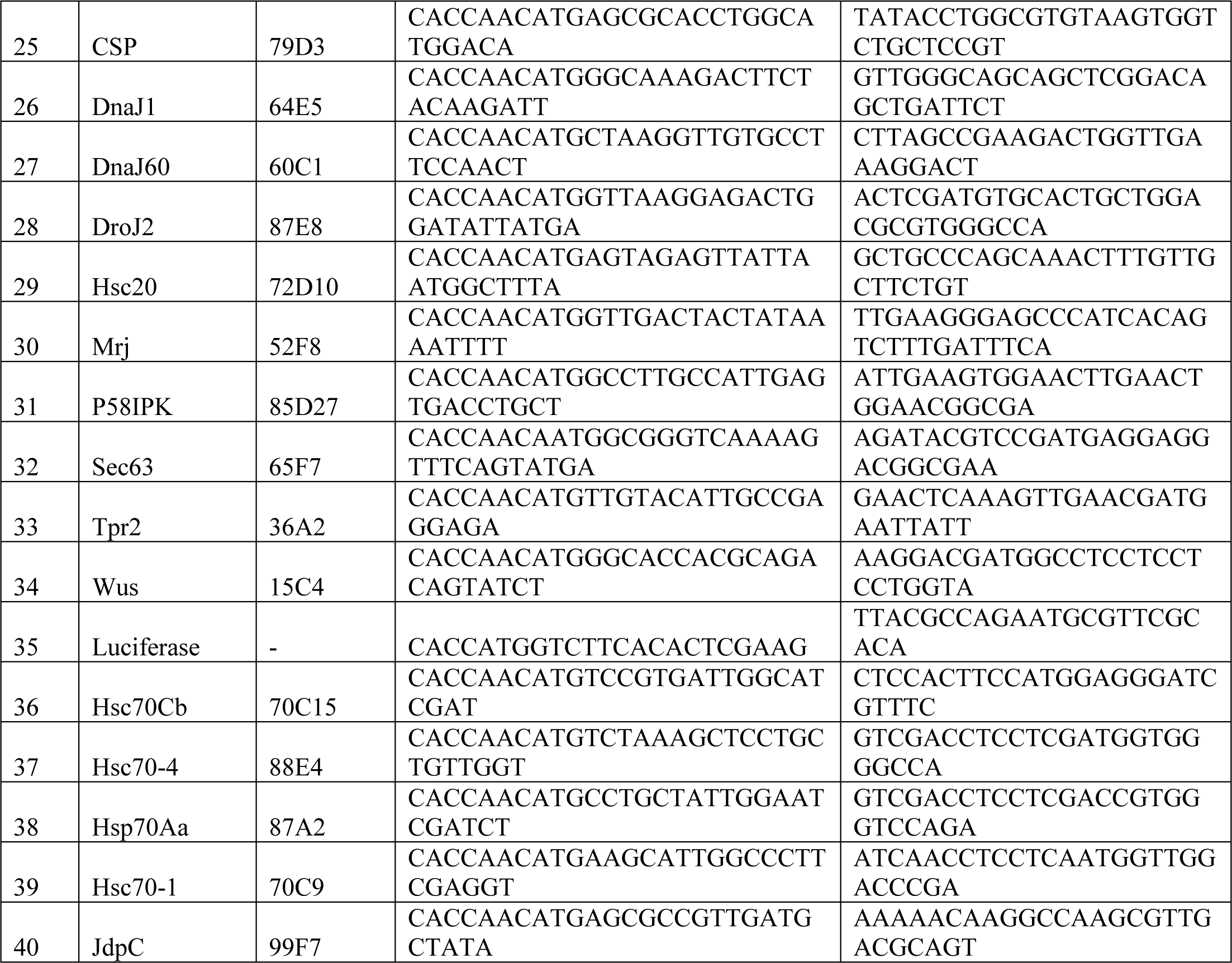
List of primers used for different DnaJ domain chaperones and co-chaperones used in Topo-D-Entr cloning.

## Notes

### Competing Interest Statement

The authors have declared no competing interest.

